# Estimation of Cross-Species Introgression Rates using Genomic Data Despite Model Unidentifiability

**DOI:** 10.1101/2021.08.14.456331

**Authors:** Ziheng Yang, Tomáš Flouri

## Abstract

Full likelihood implementations of the multispecies coalescent with introgression (MSci) model takes the genealogical fluctuation across the genome as a major source of information to infer the history of species divergence and gene flow using multilocus sequence data. However, MSci models are known to have unidentifiability issues, whereby different models or parameters make the same predictions about the data and cannot be distinguished by the data. Previous studies have focused on heuristic methods based on gene trees, and does not make an efficient use of the information in the data. Here we study the unidentifiability of MSci models under the full likelihood methods. We characterize the unidentifiability of the bidirectional introgression (BDI) model, which assumes that gene flow occurs in both directions. We derive simple rules for arbitrary BDI models, which create unidentifiability of the label-switching type. In general, an MSci model with *k* BDI events has 2^*k*^ unidentifiable modes or towers in the posterior, with each BDI event between sister species creating within-model parameter unidentifiability and each BDI event between non-sister species creating between-model unidentifiability. We develop novel algorithms for processing Markov chain Monte Carlo (MCMC) samples to remove label-switching problems and implement them in the BPP program. We analyze real and synthetic data to illustrate the utility of the BDI models and the new algorithms. We discuss the unidentifiability of heuristic methods and provide guidelines for the use of MSci models to infer gene flow using genomic data.

## INTRODUCTION

Genomic sequences sampled from modern species contain rich historical information concerning species divergences and cross-species gene flow. In the past two decades, analysis of genomic sequence data has demonstrated the widespread nature of cross-species hybridization or introgression (Baack and Rieseberg, 2007; Harrison and Larson, 2014; Mallet *et al*., 2016). A number of statistical methods have been developed to infer introgression using genomic data, most of which use data summaries such as the estimated gene trees or genome-wide site-pattern counts (Degnan, 2018; Elworth *et al*., 2019; Jiao *et al*., 2021). Full-likelihood methods applied directly to multi-locus sequence alignments (Wen and Nakhleh, 2018; Zhang *et al*., 2018; Flouri *et al*., 2020) allow estimation of evolutionary parameters including the timing and strength of introgression, as well as species divergence times and population sizes for modern and extinct ancestral species. These have moved the field beyond simply testing for the presence of cross-species gene flow.

Models of cross-species introgression are known to cause unidentifiability issues, whereby different introgression models make the same probabilistic predictions about the data, and cannot be distinguished by the data (Yu *et al*., 2012; Pardi and Scornavacca, 2015; Zhu and Degnan, 2017; Solis-Lemus *et al*., 2020). If the probability distributions of the data are identical under model *m* with parameters Θ and under model *m*′ with parameters Θ′, with

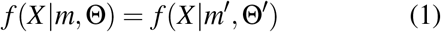

for essentially all possible data *X*, the models are unidentifiable by data *X*. Here we use the term *within-model unidentifiability* if *m* = *m*′ and Θ ≠ Θ′, or *cross-model unidentifiability* if *m ≠ m*′. In the former case, two sets of parameter values in the same model are unidentifiable, whereas in the latter, two distinct models are unidentifiable. In Bayesian inference, the prior *f*(*m*, Θ) may be used to favour a particular model or set of parameters. However, if the prior is only vaguely informative and the posterior is dominated by the likelihood, there will be multiple modes in the posterior that are not perfectly symmetrical.

Several studies examined the unidentifiability of introgression models when gene tree topologies (either rooted or unrooted) are used as data (Pardi and Scornavacca, 2015; Zhu and Degnan, 2017; Solis-Lemus *et al*., 2020), and the results apply to heuristic methods based on (reconstructed) gene trees. The issue has not been studied when full-likelihood methods are applied, which operate on multilocus sequence alignments directly. Note that unidentifiability depends on the data and the method of analysis. An introgression model unidentifiable given gene tree topologies alone may be identifiable given gene trees with coalescent times Zhu and Degnan (2017). Similarly, a model unidentifiable using heuristic methods may be identifiable when full likelihood methods are applied to the same data. It is thus important to study the problem when full likelihood methods are applied, because unidentifiability by a heuristic method may reflect its inefficient use of information in the data while unidentifiability by full likelihood methods reflects the intrinsic difficulty of the inference problem (Zhu and Yang, 2021).

Here we focus on models of episodic introgression that assume that gene flow occurs between species at fixed time points (Wen and Nakhleh, 2018; Zhang *et al*., 2018; Flouri *et al*., 2020). These are known as multispecies coalescent with introgression model (MSci; Flouri *et al*., 2020), hybrid species phylogenies (Kubatko, 2009), network multispecies coalescent model (NMSC; Zhu and Degnan, 2017), or multispecies network coalescent model (MSNC; Wen and Nakhleh, 2018; Zhang *et al*., 2018). Another class of models of between-species gene flow is the continuous migration model which assumes that migration occurs at a certain rate every generation. This is known as the isolation-with-migration (IM; Hey and Nielsen, 2004; Hey *et al*., 2018; Zhu and Yang, 2012; Dalquen *et al*., 2017) or multispecies coalescent with migration (MSC+M; Jiao *et al*., 2021) models. The IM model has very different properties concerning identifiability and is not dealt with in this study.

The bulk of the paper concerns the bidirectional-introgression (BDI) model (fig. 1), which was noted to have an unidentifiability issue (Flouri *et al*., 2020). The BDI model posits that two species coming into contact at a certain time in the past exchange genes, while the other MSci models assume introgression only in one direction. Whether gene flow tends to occur in one direction or in both directions is an interesting empirical question that may be answered by real data analyses. Here we note that recent analyses of genomic data from North-American horned lizards (Finger *et al*., 2021), the erato-sara group of *Heliconius* butterflies (Thawornwattana *et al*., 2021), and North-American chipmunks (Ji *et al*., 2021) have identified BDI events, both between sister species and between nonsister species (see also an example later). In the *Anopheles gambiae* group of African mosquitoes, introgressions between *A. gambiae* and *A. arabiensis* in both directions were suspected, but detailed analyses detected gene flow from *A. arabiensis* to *A. gambiae* only but not in the opposite direction (Thawornwattana *et al*., 2018). At any rate, BDI is one of the most plausible introgression models and appears to be one of the most common forms of cross-species gene flow. The unidentifiability of MSci models with unidirectional introgression (UDI) is simpler, and we defer its discussion to the Discussion section. Similarly we discuss unidentifiability of heuristic methods later.

**Figure 1:**
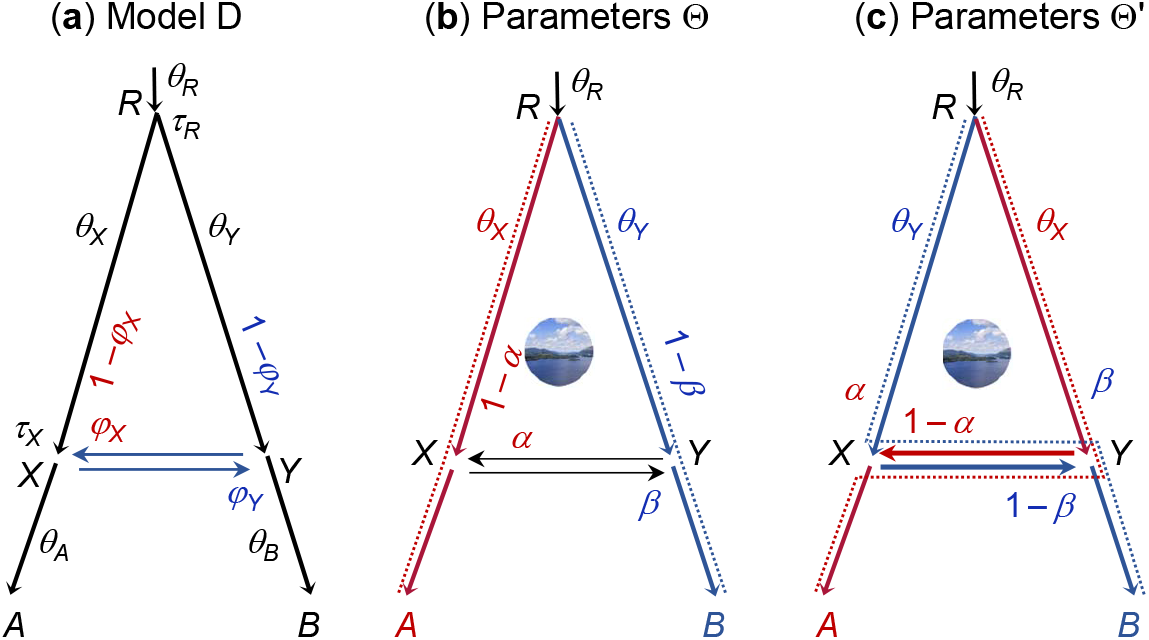
(**a**) Species tree or MSci model for two species (*A* and *B*) with a bidirectional introgression at time *τ*_*X*_ = *τ*_*Y*_, identifying nine parameters in the model. We refer to a branch by its daughter node, so that branch *XA* is also branch *A* and is assigned the population size parameter *θ*_*A*_. Both species divergence/introgression times (*τ*s) and population sizes (*θ*s) are measured in the expected number of mutations per site. (**b**) and (**c**) Two sets of unidentifiable parameters Θ and Θ′, with 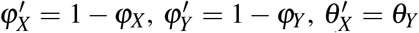, and 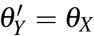, while the other five parameters (*τ*_*R*_, *τ*_*X*_ = *τ*_*Y*_, *θ*_*A*_, *θ*_*B*_, and *θ*_*R*_) are identical between Θ and Θ′. The dotted lines indicate the main routes taken by sequences sampled from species *A* and *B*, if both introgression probabilities *α* and *β* are 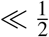.

The basic BDI model between two species (fig. 1) involves nine parameters, with Θ = (*θ*_*A*_, *θ*_*B*_, *θ*_*X*_, *θ*_*Y*_, *θ*_*R*_, *τ*_*R*_, *τ*_*X*_, *φ*_*X*_, *φ*_*Y*_). An introgression model is similar to a species tree except that it includes horizontal branches representing lateral gene flow across species. Besides speciation nodes representing species divergences, there are hybridization nodes representing introgression events as well. While a speciation node has one parent and two daughters, a hybridization node has two parents and one daughter. The ‘introgression probabilities’ or ‘admixture proportions’ (*φ* and 1 − *φ*) specify the contributions of the two parental populations to the hybrid species. When we trace the genealogical history of a sample of sequences from the modern species backwards in time and reach a hybridization node, each of the sequences takes the two parental paths with probabilities *φ* and 1 − *φ*. There are thus three types of parameters in an introgression (or MSci) model: the times of species divergence and introgression (*τ*s), the (effective) population sizes of modern and ancestral species (*θ*s), and the introgression probabilities (*φ*s). Both the divergence times (*τ*s) and population sizes (*θ*s) are measured in the expected number of mutations per site.

The BDI model, in the case of two species (fig. 1), is noted to have an unidentifiability issue (Flouri *et al*., 2020). Let Θ′ be a set of parameters with the same values as Θ except that 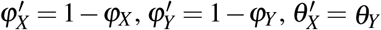, and 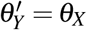. Then *f*(*G*|Θ) = *f*(*G*|Θ′) for any gene tree *G* (fig. 1b&c). Here *G* represents both the gene tree topology and branch lengths (coalescent times). We assume multiple sequences sampled per species at the same locus (see Discussion for unidentifiability caused by sampling only one sequence per species). Thus for every point Θ in the parameter space, there is a ‘mirror’ point Θ′ with exactly the same likelihood. With Θ, the *A* sequences take the left (upper) path at *X* and enter population *RX* with probability 1 − *φ*_*X*_, coalescing at the rate 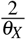, while with Θ′, the same *A* sequences may take the right (horizontal) path and enter population *RY* with probability 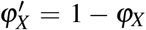, coalescing at the rate 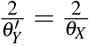. The differences between Θ and Θ′ are in the labelling, with ‘left’ and *X* under Θ corresponding to ‘right’ and *Y* under Θ′, but the probabilities involved are the same. The same argument applies to sequences from *B* going through node *Y*, and to any numbers of sequences from *A* and *B* considered jointly. Thus *f*(*G*|Θ) = *f*(*G*|Θ′) for essentially all *G*. If the priors on *φ*_*X*_ and *φ*_*Y*_ are symmetrical, say *φ*∼ beta(*α, α*), the posterior density will satisfy *f* (Θ | *X*) = *f* (Θ′ |*X*) for all *X*. Otherwise the “twin towers” in the posterior may not have exactly the same height.

The situation is very similar to the label-switching problem in Bayesian clustering (Richardson and Green, 1997; Celeux *et al*., 1998; Stephens, 2000; Jasra *et al*., 2005). Consider data *X* = {*x*_*i*_} as a sample from a mixture of two normal distributions, ℕ(*μ*_1_, 1) and ℕ(*μ*_2_, 1), with the mixing proportions *p*_1_ and *p*_2_ = 1 − *p*_1_. Let Θ = (*p*_1_, *μ*_1_, *μ*_2_) be the parameter vector. Then Θ′ = (*p*_2_, *μ*_2_, *μ*_1_) with *p*_2_ = 1 − *p*_1_ will have exactly the same likelihood, so that *f*(*X* |Θ) = *f*(*X*| Θ′) for essentially all data *X*. In effect, the labels ‘group 1’ and ‘group 2’ are switched between Θ and Θ′.

As an example, we fit the BDI model of figure 2a to the first 500 noncoding loci on chromosome 1 in the genomic data from three *Heliconius* butterfly species: *H. melpomene, H. timareta*, and *H. numata* (Edelman *et al*., 2019; Thawornwattana *et al*., 2021). Figure 3a shows the trace plots for parameters *φ*_*X*_ and *φ*_*Y*_ from a Markov chain Monte Carlo (MCMC) run. The Markov chain moves between two peaks, centered around (*φ*_*X*_, *φ*_*Y*_) = (0.35, 0.1) and (0.65, 0.9), respectively. In effect, the algorithm is switching between Θ and Θ′ and changing the definition of parameters during the same MCMC run. This is a label-switching problem, as occurs in Bayesian clustering. The usual practice of estimating parameters by their posterior means calculated using the MCMC sample (0.54 for *φ*_*X*_ and 0.62 for *φ*_*Y*_ in fig. 3a) and constructing the credibility intervals is inappropriate. Indeed the posterior distribution of Θ is exactly symmetrical with twin towers, and if the chain is run long enough, the sample means of *φ*_*X*_ and *φ*_*Y*_ will be exactly 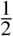, irrespectively of what values may fit the data well. The results are similar when the first 500 exonic loci are analyzed, in which the Markov chain moves between two towers centered around (0.3, 0.1) and (0.7, 0.9) (fig. S1a).

**Figure 2:**
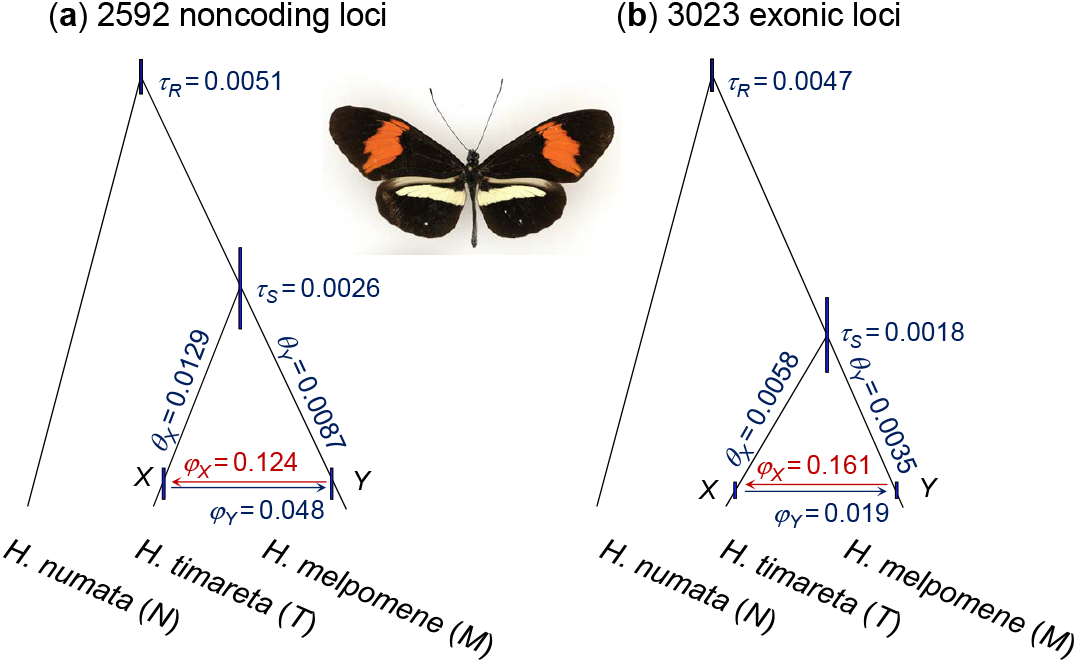
Species tree or BDI model for *Heliconius melpomene, H. timareta*, and *H. numata*. The branches are drawn to represent the posterior means of divergence/introgression times obtained from BPP analysis of (**a**) the 2592 noncoding and (**b**) the 3023 exonic loci from chromosome 1, while the node bars represent the 95% HPD CIs. See table 1 for estimates of all parameters. Photo of *H. timareta* courtesy of James Mallet.

**Figure 3:**
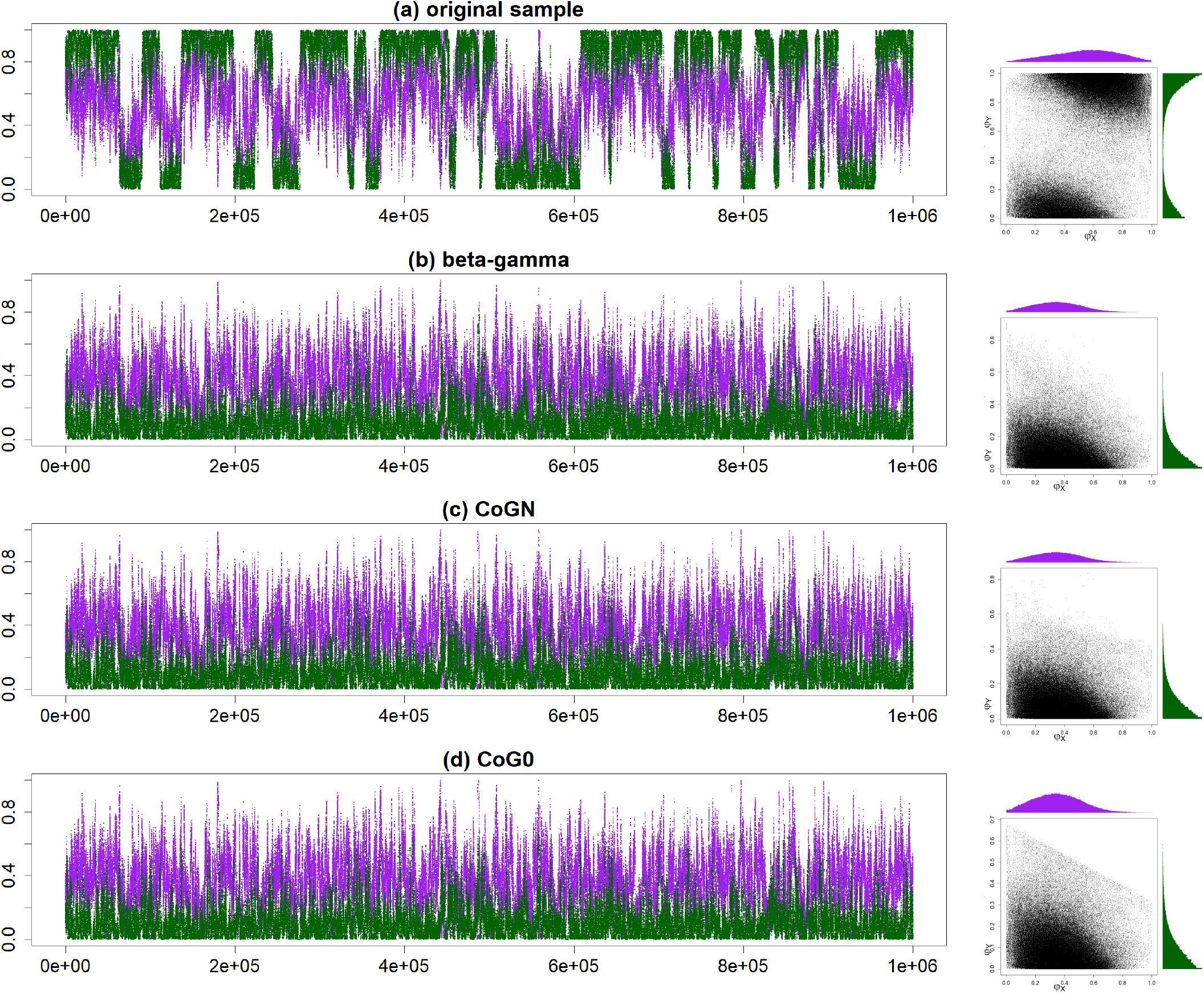
Trace plots of MCMC samples and 2-D scatter plots for parameters *φ*_*X*_ (purple) and *φ*_*Y*_ (green) (**a**) before and (**b**–**d**) after the post-processing of the MCMC sample in the BPP analysis of the first 500 noncoding loci from chromosome 1 of the *Heliconius* data under the MSci model of figure 2. The three algorithms used are (**b**) *β*−*γ*, (**c**) CoG_*N*_, and (**d**) CoG_0_.

Unidentifiable models cannot be applied to real data as they are trying to “distinguish the indistinguishable” (Pardi and Scornavacca, 2015). Results such as those of figures 3a & S1a raise two questions. First, what are the rules concerning the unidentifiability of general BDI models, for example, if there are more than two species on the species tree, more than one BDI event, or if the BDI event involves non-sister species (fig. 1). Second how do we deal with the problem of label-switching and make the models useful for real data analyses? We address those problems. We study the unidentifiability issue of BDI models for an arbitrary number of species with an arbitrary species tree, when a full-likelihood method is applied to multilocus sequence data. It has been conjectured that an MSci model is identifiable by full likelihood methods using data of multi-locus sequence alignments if and only if it is identifiable when the data consist of gene trees with coalescent times (Flouri *et al*., 2020). We make use of this conjecture and consider two BDI models to be unidentifiable if and only if they generate the same distribution of gene trees with coalescent times. We emphasize that the unidentifiability discussed here affects all methods of inference using genomic sequence data, including heuristic methods based on summary statistics as well as full likelihood methods (see Discussion). We identify general rules for the unidentifiability of the BDI models. We then develop new algorithms for post-processing the MCMC samples generated from a Bayesian analysis under the BDI model to remove the label-switching. Those efforts make the BDI models usable for real data analysis despite their unidentifiability. We use the BPP program (Flouri *et al*., 2018) to analyze synthetic datasets as well as genomic data from *Heliconius* butterflies to demonstrate the utility of the BDI models and the new algorithms. After we have dealt with the BDI models, we discuss the unidentifiability of UDI models and of heuristic methods.

**Table 1.** Posterior means and 95% HPD CIs (in parenthees) for parameters in the BDI model of figure 2 for the *Heliconius* data.

| | Noncoding, $L = 500$ | Noncoding, $L = 2592$ | Exonic, $L = 500$ | Exonic, $L = 3023$ |
| --- | --- | --- | --- | --- |
| $\tau_R$ | 4.73 (4.33, 5.13) | 5.10 (4.89, 5.30) | 4.39 (3.98, 4.81) | 4.71 (4.54, 4.88) |
| $\tau_S$ | 3.12 (2.05, 4.19) | 2.58 (2.12, 3.05) | 1.95 (1.07, 2.82) | 1.78 (1.38, 2.19) |
| $\tau_X = \tau_Y$ | 0.62 (0.21, 1.02) | 0.25 (0.09, 0.40) | 0.20 (0.03, 0.37) | 0.13 (0.05, 0.24) |
| $\theta_M$ | 1.50 (0.62, 2.34) | 0.69 (0.35, 1.10) | 0.38 (0.08, 0.70) | 0.32 (0.14, 0.52) |
| $\theta_T$ | 2.55 (1.40, 3.74) | 1.23 (0.65, 1.84) | 0.79 (0.13, 1.28) | 0.63 (0.32, 0.94) |
| $\theta_N$ | 15.1 (12.0, 18.5) | 23.0 (20.3, 25.7) | 11.2 (9.11, 13.5) | 12.4 (11.4, 13.4) |
| $\theta_R$ | 5.08 (4.12, 6.05) | 5.74 (5.23, 6.24) | 5.76 (4.83, 6.70) | 6.68 (6.24, 7.11) |
| $\theta_S$ | 4.62 (1.85, 7.40) | 6.92 (5.48, 8.37) | 5.31 (3.38, 7.36) | 7.50 (6.51, 8.49) |
| $\theta_X$ | 11.40 (2.83, 21.2) | 12.90 (7.35, 19.6) | 8.04 (1.67, 15.4) | 5.80 (3.60, 8.36) |
| $\theta_Y$ | 6.78 (2.42, 11.6) | 8.74 (5.69, 12.0) | 4.03 (0.60, 7.51) | 3.49 (2.56, 4.50) |
| $\phi_X$ | 0.354 (0.022, 0.664) | 0.124 (0.007, 0.243) | 0.280 (0.002, 0.547) | 0.161 (0.070, 0.264) |
| $\phi_Y$ | 0.104 (0.000, 0.306) | 0.048 (0.000, 0.139) | 0.116 (0.000, 0.318) | 0.019 (0.000, 0.056) |

Note.— Estimates of $\tau$ s and $\theta$ s are multiplied by $10^3$ . MCMC samples are processed using the $\beta$ - $\gamma$ algorithm before they are summarized.

## THEORY

### The rule of unidentifiability of BDI models

In full likelihood implementations of the MSci model, the gene tree *G* for any given sample of sequences from the modern species represents the complete history of coalescence and introgression events for the sample, including the gene tree topology, the coalescent times, as well as the parental path taken by each sequence at each hybridization node (e.g., Jiao *et al*., 2021, eq. 14). The probability distribution of the gene tree *G* depends on the species tree, species divergence times (*τ*s), the population sizes (*θ*s) which determine the coalescent rates in the different populations 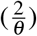, and the introgression probabilities at the hybridization nodes (*φ*). It does not depend on the labels attached to the internal nodes in the species tree.

Consider a part of the species tree or MSci model where species *A* and *B* exchange migrants at time *τ*_*X*_ = *τ*_*Y*_ (fig. 4). To study the backwards-in-time process of coalescent and introgression, which gives the probability density of the gene tree *f*(*G* |*S*, Θ), we can treat nodes *X* and *Y* as one node, *XY*. When sequences from *A* reach node *XY*, each of them has probability 1 − *φ*_*X*_ of taking the left parental path (*RX*) and probability *φ*_*X*_ of taking the right parental path (*SY*). Similarly when sequences from *B* reach node *XY*, they have probabilities *φ*_*Y*_ and 1 − *φ*_*Y*_ of taking the left (*RX*) and right (*YS*) parental paths, respectively. If we swap branches *A* and *B*, carrying with them their population size parameters (*θ*) and introgression probabilities (*φ*), the probability density of the gene-trees remains unchanged. Thus the species tree-parameter combinations (*S*, Θ) and (*S*′, Θ′) of figure 4b&c give exactly the same probability distribution,

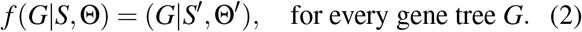

**Figure 4:**
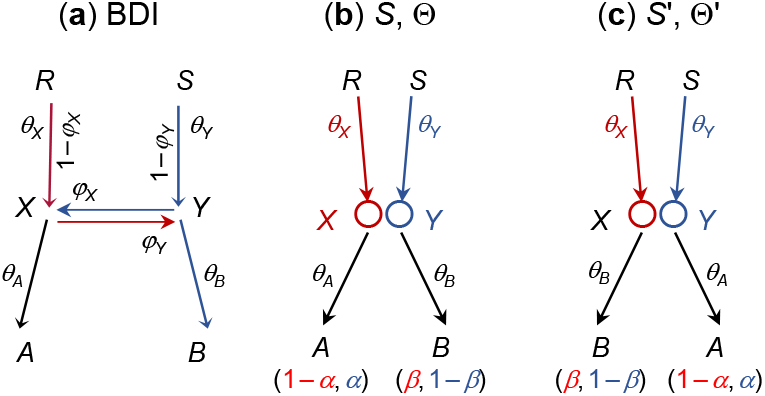
A part of a species tree (MSci model) for illustrating the rule of BDI unidentifiability. (**a**) In the BDI model, species *RXA* and *SYB* exchange migrants at time *τ*_*X*_ = *τ*_*Y*_. Treat *X* and *Y* as one node with left parent *RX* with population size *θ*_*X*_ and right parent *SY* with population size *θ*_*Y*_. When a sequence from *A* reaches *XY*, it takes the left and right parental paths with probabilities 1 − *φ*_*X*_ and *φ*_*X*_, respectively. When a sequence from *B* reaches *XY*, it goes left and right with probabilities *φ*_*Y*_ and 1 − *φ*_*Y*_, respectively. (**b & c**) Placing the two daughters in the order (*A, B*) as in Θ or (*B, A*) as in Θ′ does not affect the distribution of gene trees, and constitutes unidentifiable towers in the posterior space. If *X* and *Y* are sister species and have the same mother node (with *R* and *S* to be the same node), the unidentifiability is within-model; otherwise it is cross-model.

In other words, (*S*, Θ) and (*S*′, Θ′) are unidentifiable (eq. 1).

Note that the processes of coalescent and introgression before reaching nodes *A* and *B* (with time running backwards) are identical between Θ and Θ′, as are the processes past nodes *X* and *Y*. Thus the rule applies if each of *A* and *B* is a subtree, with introgression events inside, or if there are introgression events involving a descendant of *A* and a descendant of *B*.

If *A* and *B* are sister species or the parents *R* and *S* are one node in the species tree, the species trees (*A, B*) and (*B, A*) will be the same so that *S* = *S*′ in eq. 2. Then Θ and Θ′ (fig. 4) will be two sets of parameter values in the same model and we have a case of within-model unidentifiability. Otherwise the unidentifiability is cross-model.

### Canonical cases of BDI models

Here we study major BDI models to illustrate the rule of unidentifiability and to provide reference for researchers who may apply those models to analyze genomic datasets.

If we add subtrees onto branches *XA, YB*, or the root branch *R* in the two-species tree of figure 1a, so that the BDI event remains to be between two sister species, the model will exhibit within-model parameter unidentifiability (fig. S2), just like the basic model of figure 1a.

If the BDI event is between non-sister species, the model exhibits cross-model unidentifiability. Figures S3a&a′ show a model with a BDI event between cousins, while in figures S3b&b′, the two species involved in the BDI event are more distantly related.

Figures S4a, b &c show three models each with a BDI event between non-sister species. In figure S4a, *X* and *Y* are non-sister species on the original binary species tree. In figure S4b&c, *X* and *Y* are non-sister species because there are introgression events involving branches *RX* and/or *RY*. In all three cases, there is cross-model unidentifiability, with the twin towers shown in S4a′, b′&c′.

The case of two non-sister BDI events for three species is illustrated in figure S5. According to our rule, there are four unidentifiable models in the posterior, with parameter mappings shown in figure S5. One way of seeing that the four models are equivalent or unidentifiable is to assume that the introgression probabilities (*φ*_*X*_, *φ*_*Y*_, *φ*_*Z*_, and *φ*_*W*_) are all 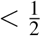, and then work out the major routes taken when we trace the genealogical history of sequences sampled from modern species. In such cases, all four models of figure S5 predict the following: most sequences from *A* will take the route *ZR* at node *ZW* with probability 1 − *γ*; most sequences from *B* will take the route *X* -*W* at node *XY* (with probability 1 − *α*), then the route *WS* at node *ZW* (with probability 1 − *δ*), before reaching *SR*; and most sequences from *C* will take the route *YS* at node *XY* (with probability 1 − *β*), before reaching *SR*. Of course the four models are unidentifiable whatever values the introgression probabilities take. Those models have been used to analyze genomic data from Texas Horned Lizards (*Phrynosoma cornutum*) (Finger *et al*., 2021).

Figure 5 shows two models for five species, each model involving three BDI events. In figure 5a, all three BDI events involve sister species, so that there are 2^3^ = 8 unidentifiable within-model towers in the posterior. In figure 5b, one BDI event involves non-sister species while two involve sister species, so that there are two unidentifiable models, each of which has four unidentifiable within-model towers in the posterior.

**Figure 5:**
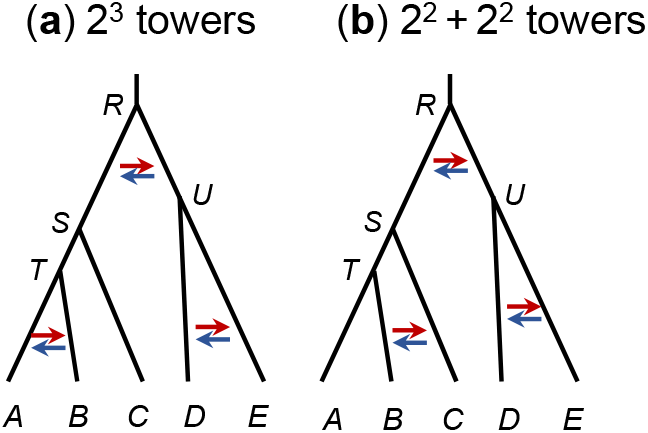
Two species trees (MSci models) for five species each with three BDI events. (**a**) Three BDI events between sister species create 2^3^ = 8 within-model towers in the posterior. (**b**) Two BDI events between sister species and one BDI event between non-sister species create two unidentifiable models each with four within-model unidentifiable towers in the posterior space.

In general, if there are *m* BDI events between sister species and *n* BDI events between non-sister species, there will be 2^*n*^ unidentifiable models, each having 2^*m*^ within-model unidentifiable towers.

### Unidentifiability of double-DBI models

Figure 6a shows two BDI events between species *A* and *B*, which occurred at times *τ*_*X*_ = *τ*_*Y*_ and *τ*_*Z*_ = *τ*_*W*_, respectively. To apply the rule of figure 4, we treat *Z* and *W* as one node so that *X* and *Y* are considered sister species. There are then four within-model unidentifiable towers in the posterior space, shown as Θ_1_-Θ_4_ in fig. 6. The parameter mappings are given in the following table

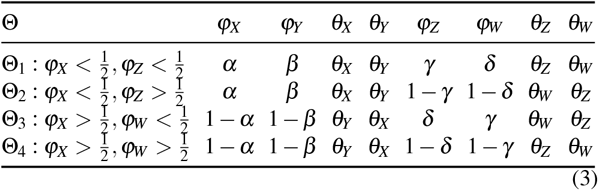

**Figure 6:**
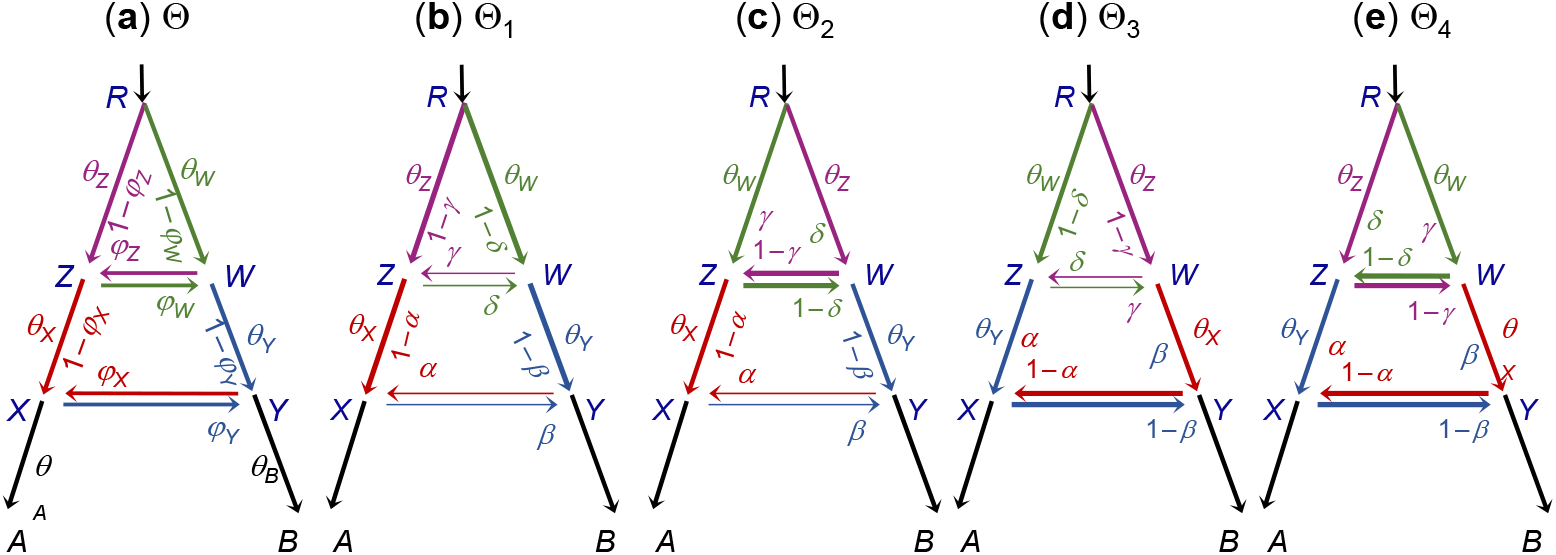
Species trees (MSci models) for two species (*A* and *B*) with double DBI events creating four within-model towers, represented by Θ_1_, Θ_2_, Θ_3_, and Θ_4_. (**a**) The model involves 14 parameters: 7 *θ*s, 3 *τ*s, and 4 *φ*s, with eight of them involved in the label-switching unidentifiability, Θ = (*φ*_*X*_, *φ*_*Y*_, *θ*_*X*_, *θ*_*Y*_, *φ*_*Z*_, *φ*_*W*_, *θ*_*Z*_, *θ*_*W*_). (**b**)-(**e**) Four unidentifiable towers showing the mappings of parameters (eq. 3). To apply the rule of figure 4, we treat each pair of BDI nodes as one node, so that *X* and *Y* have the same node *ZW* as the parent, and the unidentifiability caused by the BDI event at nodes *X* -*Y* is within-model, as is the unidentifiability for the BDI event at nodes *Z*-*W*.

In general, with *k* BDI events between two species, which occurred at different time points in the past, there will be 2^*k*^ unidentifiable within-model towers in the posterior. There may be little information in practical datasets to estimate so many parameters: if all sequences have coalesced before they reach the ancient introgression events near the root of the species tree, the introgression probabilities (*φ*s) and the associated population sizes (*θ*s) will be nearly impossible to estimate. Thus we do not consider more than two BDI events between two species. Note that even the model with one BDI event is not identifiable by heuristic methods that use gene tree topologies only. A small simulation is conducted to illustrate the feasibility of applying the double-BDI model (fig. 6) to genomic datasets; see Results.

### Addressing unidentifiability issues and difficulties with identifiability constraints

According to our rule, MSci models with BDI events can create both within-model and cross-model unidentifiability. Cross-model unidentifiability is relatively simple to identify and deal with. If the MCMC is run with the MSci model fixed (Flouri *et al*., 2020), only one of the models (e.g., model *S*_1_ with parameters Θ_1_ in fig. S5) is visited in the chain. One can then summarize the posterior distribution for parameters under that model (which may be smooth and single-moded), and the posterior summary may be mapped onto the other unidentifiable models according to the rule. See Finger *et al*. (2021) for such an application of BDI models of figure S5. If the MCMC is trans-model and visits different models in the chain (Zhang *et al*., 2018; Wen and Nakhleh, 2018), the posterior space is symmetrical between the unidentifiable models (such as models *S*_1_– *S*_4_ of fig. S5). However, such symmetry is unlikely to be achieved in the MCMC sample due to well-known mixing difficulties of trans-model MCMC algorithms. One may choose to focus on one of the models (e.g., *S*_1_ of fig. S5) and post-process the MCMC sample to map the sample onto the chosen model before producing the within-model posterior summary. Oftentimes the MCMC may explore the within-model posterior space very well, despite difficulties of moving from one model to another. In all cases, the researcher has to be aware of the unidentifiable models which are equally good explanations of the data (see Discussion).

Our focus here is on within-model unidentifiability created by BDI events between sister species. When there are multiple modes in the posterior, each mode may offer a sensible interpretation of the data, but it is inappropriate to merge MCMC samples from different modes, or to construct posterior summaries such as the posterior means and CIs using MCMC samples that traverse different modes. It is instead more appropriate to summarize the samples for each mode.

A common strategy for removing label-switching is to apply so-called *identifiability constraints*. In the simple BDI model of figure 1, any of the following constraints may be applicable: 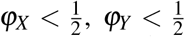, and *θ*_*X*_ *< θ*_*Y*_. Such identifiability constraints may be imposed during the MCMC or during post-processing of the MCMC samples. As discussed previously (Celeux *et al*., 1998; Stephens, 2000), such a constraint may be adequate if the posterior modes are well separated, but may not work well otherwise. For example, if *φ*_*X*_ is far away from 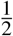 in all MCMC samples, it will be simple to post-process the MCMC sample to impose the constraint 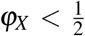. This is the case in analyses of the large datasets in this paper, for example, when all noncoding and exonic loci from chromosome 1 of the *Heliconius* data are analyzed (table 1). However, when the posterior modes are not well-separated (either because the true parameter value is close to the boundary defined by the inequality or because the data lack information so that the CIs are wide), different identifiability constraints can lead to very different parameter posteriors (Richardson and Green, 1997), and an appropriate constraint may not exist. A serious problem in such cases is that imposing an identifiability constraint may generate posterior distributions over-represented near the boundary, with seriously biased posterior means (Celeux *et al*., 1998; Stephens, 2000). For example, *φ*_*X*_ may have substantial density mass both below and above 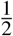, and imposing the constraint 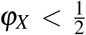 will artificially generate high density mass close to 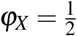. Similarly the posterior distributions of *θ*_*X*_ and *θ*_*Y*_ may overlap, so that the constraint *θ*_*X*_ *< θ*_*Y*_ may not be appropriate.

### New algorithms to process MCMC samples from the BDI model to remove label switching

One approach to dealing with label-switching problems in Bayesian clustering is *relabelling*. The MCMC is run without any constraint, and the MCMC sample is then post-processed to remove label switching, by attempting to move each point in the MCMC sample to its alternative unidentifiable positions in order to, as far as possible, make the marginal posterior distributions smooth and unimodal (Celeux *et al*., 1998; Stephens, 2000). The processed sample is then summarized to generate the posterior of the parameters. Here we follow this strategy and implement three relabelling algorithms to post-process the MCMC samples generated under the BDI model.

Let Θ = (*φ*_*X*_, *φ*_*Y*_, *θ*_*X*_, *θ*_*Y*_), which has a mirror point 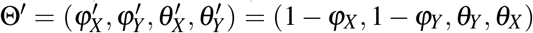 (fig. 1). The other parameters in the model are not involved in the unidentifiability and are simply copied along. Let Θ_*t*_, *t* = 1, · · ·, *N*, be the *N* samples of parameters generated by the MCMC algorithm. Each sample is a point in the 4-D space. Let *z*_*t*_ be a transform for point *t*, with *z*_*t*_(Θ_*t*_) = Θ_*t*_ to be the original point, and *z*_*t*_(Θ_*t*_) = Θ′ to be the transformed or mirror point (fig. 1b&c). With a slight abuse of notation, we also treat *z*_*t*_ as an indicator, with *z*_*t*_ = 0 and 1 representing Θ_*t*_ and 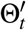, respectively. For each sample *t*, we choose either the original point or its mirror point, to make the posterior of the parameters look smooth and singlemoded as far as possible. The first two algorithms, called center-of-gravity algorithms CoG_0_ and CoG_*N*_, loop through two steps.

### Algorithms CoG_0_ and CoG_*N*_

Initialize. For each point *t, t* = 1, · · ·, *N*, pick either the original point (Θ_*t*_) or its mirror point 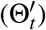. We set *z*_*t*_ to 0 (for the original point Θ_*t*_) if *φ*_*X*_ + *φ*_*Y*_ *<* 1 or 1 (for the mirror point 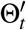) otherwise.

- Step 1. Determine the center of gravity, given by the sample means of the parameters, 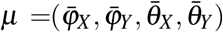.
- Step 2. For each point *t* = 1, · · ·, *N*, compare the current and its mirror positions and choose the one closer to the center of gravity (*μ*).

In step 2, we use the Euclidean distance

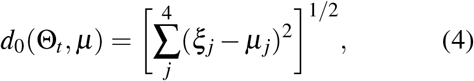

where *ξ*_*j*_ are the four parameters in Θ_*t*_: *φ*_*X*_, *φ*_*Y*_, *θ*_*X*_, *θ*_*Y*_. This is algorithm CoG_0_.

If we consider different scales in the different dimensions (for example, *φ*_*X*_ and *θ*_*X*_ may have very different posterior variances), we can calculate the sample variances *ν* (in addition to the sample means *μ*) in step 1 and use them as weights to normalize the differences in step 2, with

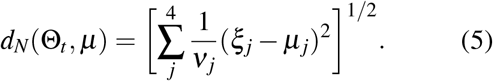

We refer to this as algorithm CoG_*N*_.

Note that each MCMC sample point Θ_*t*_ can be in either of two positions (represented by *z*_*t*_ = 0 or 1). Step 1 calculates the center of attraction (*μ*), which represents the current ‘location of most points’. Given the center of attraction, step 2 moves each point to the position closer to the center of attraction. If the posterior has two modes due to label switching but no other modes, all points will be moved to the same neighborhood around the center of attraction. Which of the two unidentifiable modes becomes the center of attraction is arbitrary, influenced by the initial positions when the algorithm runs.

The third algorithm, called the *β*−*γ* algorithm, follows the relabelling algorithm for Bayesian clustering of Stephens (2000). We use maximum likelihood (ML) to fit the sample {Θ_*t*_} to independent beta distributions for *φ*_*X*_ and *φ*_*Y*_ and gamma distributions for *θ*_*X*_ and *θ*_*Y*_ :

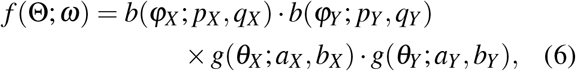

where

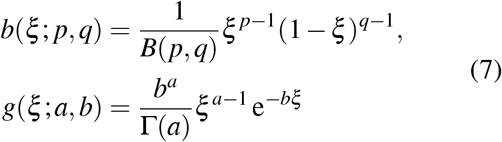

are the beta and gamma densities and where *ω* = (*p*_*X*_, *q*_*X*_, *p*_*Y*_, *q*_*Y*_, *a*_*X*_, *b*_*X*_, *a*_*Y*_, *b*_*Y*_) is the vector of parameters in those densities.

The log likelihood, as a function of the parameters *ω* and the transforms *z* = {*z*_*t*_}, is

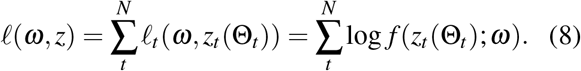

We have implemented the following iterative algorithm to estimate *ω* and *z* by maximizing *ℓ*.

### Algorithm *β*−*γ*

Initialize *z*_*t*_, *t* = 1, · · ·, *N*. As before, we set *z*_*t*_ to 0 (for Θ_*t*_) if *φ*_*X*_ + *φ*_*Y*_ *<* 1 or 1 (for 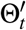) otherwise.

- Step 1. Choose 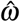 to maximize the log likelihood *ℓ* (eq. 8) with the transforms *z* fixed.
- Step 2. For *t* = 1, · · ·, *N*, choose *z*_*t*_ = 0 or 1 to maximize 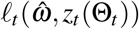 with 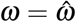 fixed. In other words compare Θ_*t*_ and 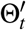 and choose the one that better fits the beta and gamma distributions.

Step 1 fits two beta and two gamma distributions by ML and involves four separate 2-D optimization problems. The maximum likelihood estimates (MLEs) of *p* and *q* for the beta distribution *b*(*ξ* ; *p, q*) are functions of Σ_*t*_ log *ξ*_*t*_ and Σ_*t*_ log(1 − *ξ*_*t*_), whereas the MLEs of *a* and *b* for the gamma distribution *g*(*ξ* ; *a, b*) are functions of Σ_*t*_ *ξ*_*t*_ and Σ_*t*_ log *ξ*_*t*_. These optimization problems are simple, which we solve using the BFGS algorithm in the paml program (Yang, 2007). Step 2 involves *N* independent optimization problems, each comparing two points (*z*_*t*_ = 0 and 1), with *ω* fixed. It is easy to see that the algorithm is nondecreasing (that is, the log likelihood *ℓ* never decreases) and converges, as step 1 involves ML estimation of parameters in the beta and gamma distributions, and step 2 involves comparing two points.

Note that the *β*−*γ* algorithm becomes the CoG_0_ and CoG_*N*_ algorithms if the beta and gamma densities are replaced by normal densities (with the same or different variances for CoG_0_ and CoG_*N*_, respectively).

For illustration we applied the CoGN_0_ algorithm to a ‘thinned’ sample of 1000 points from the MCMC sample of figure 3a generated in the BPP analysis of the 500 noncoding *Heliconius* loci. We used three initial conditions (three rows in fig. S6). The last plot on each row is a summary of the final processed sample. Thus the first two runs produced the same posterior, while the third run produced its mirror image. Note that the relabelling algorithms remove the label-switching problems, but do not remove unidentifiability.

### Algorithms CoG_0_, CoG_*N*_, and *β*−*γ* for the double-BDI model

Under the double-BDI model (fig. 6a), there are four within-model unidentifiable towers, specified by eight parameters. Thus *z*_*t*_ takes four values (0, 1, 2, 3), and Θ = (*φ*_*X*_, *φ*_*Y*_, *θ*_*X*_, *θ*_*Y*_, *φ*_*Z*_, *φ*_*W*_, *θ*_*Z*_, *θ*_*W*_). We use the same strategy and fit four beta distributions to the *φ*s and four gamma distributions to the *θ*s, so that there are 16 parameters in *ω*. We implement the three algorithms (*β*−*γ*, CoG_*N*_, and CoG_0_) as before. We prefer the tower in which the introgression probabilities are small and initialize the algorithm accordingly. The transforms (*z*_*t*_ = 0, 1, 2, 3) are as follows (eq. 3)

- *z*_*t*_ = 0: if the parameters are in Θ_1_, do nothing.
- *z*_*t*_ = 1: if in Θ_2_, let *φ*_*Z*_ = 1 − *φ*_*Z*_, *φ*_*W*_ = 1 − *φ*_*W*_, and swap *θ*_*Z*_ and *θ*_*W*_.
- *z*_*t*_ = 2: if in Θ_3_, let *φ*_*X*_ = 1 − *φ*_*X*_, *φ*_*Y*_ = 1 − *φ*_*Y*_, swap *θ*_*X*_ and *θ*_*Y*_, swap *φ*_*Z*_ and *φ*_*W*_, swap *θ*_*Z*_ and *θ*_*W*_ ;
- *z*_*t*_ = 3: if in Θ_4_, let *φ*_*X*_ = 1 − *φ*_*X*_, *φ*_*Y*_ = 1 − *φ*_*Y*_, swap *θ*_*X*_ and *θ*_*Y*_, and let *φ*_*Z*_ = 1 − *φ*_*W*_ and *φ*_*W*_ = 1 − *φ*_*Z*_.

The algorithm similarly loops through two steps. In step 1 we calculate the center of gravity (represented by the means) or estimate parameters 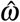 to fit the beta and gamma densities, with the transforms *z* fixed. For CoG_0_ and CoG_*N*_, this step involves calculating the sample means and variances for the eight parameters in Θ, while for *β*−*γ*, it involves a 16-D optimization problem (or eight 2-D optimization problems) for fitting the beta and gamma distributions by ML. In step 2, we compare the four positions for each sample point when the center of gravity or parameters 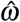 are fixed.

To apply the rule and the algorithms developed here, we need to identify the BDI event and the parameters involved in the unidentifiability, that is, (*φ*_*X*_, *φ*_*Y*_, *θ*_*X*_, *θ*_*Y*_) under the BDI model, or (*φ*_*X*_, *φ*_*Y*_, *θ*_*X*_, *θ*_*Y*_, *φ*_*Z*_, *φ*_*W*_, *θ*_*Z*_, *θ*_*W*_) under double-BDI. The algorithm is then used to process the MCMC sample. If there are multiple BDI or double-BDI events between sister species, one may simply apply the post-processing algorithm multiple times. For instance, three rounds of post-processing may be applied for the model of figure 5a (for the BDI events between *A* and *B*, between *D* and *E*, and between *S* and *U*, respectively), while the model of 5b requires two rounds (for the BDI between *D* and *E*, and between *S* and *U*).

The algorithms are implemented in C and require minimal computation and storage. Processing 5 × 10^5^ samples takes several rounds of iteration and a few seconds of running time, mostly spent on reading and writing files. The algorithms are integrated into the BPP program (Flouri *et al*., 2018) so that MCMC samples from various BDI models are post-processed and summarized automatically. We also provide a stand-alone program in the github repository `abacus-gene/bpp-msci-D-process-mcmc/`.

## RESULTS

### Introgression between *Heliconius melpomene* and *H. timareta*

We fitted the BDI model of figure 2 to the genomic sequence data from three species of *Heliconius* butterflies: *H. melpomene, H. timareta*, and *H. numata* (Edelman *et al*., 2019; Thawornwattana *et al*., 2021). When we used the first 500 loci, either noncoding or exonic, there was substantial uncertainty in the posterior of *φ*_*X*_ and *φ*_*Y*_, and the MCMC jumped between the twin towers, and the marginal posteriors had multiple modes, due to label switching (figs. 3a & S1a). Post processing of the MCMC sample using the new algorithms led to single-moded marginal posterior distributions (figs. 3b–d & S1b–d). The three algorithms produced extremely similar results for both datasets. For example, the posterior mean and 95% CI for *φ*_*X*_ from the noncoding data were 0.356 (0.026, 0.671) by CoG_0_, 0.357 (0.026, 0.674) by CoG_*N*_, and 0.354 (0.022, 0.664) by *β*−*γ*, while those for *φ*_*Y*_ were 0.103 (0.000, 0.304) by CoG_0_ and CoG_*N*_, and 0.104 (0.000, 0.306) by *β*−*γ*.

We then analyzed all the 2592 noncoding and 3023 exonic loci on chromosome 1. With the large datasets, the parameters were better estimated with narrower CIs and the unidentifiable towers were well-separated. In fact, the MCMC visited only one of the two towers, but the visited tower was well explored so that multiple runs produced highly consistent results after label-switching was removed using the relabelling algorithms. When we started the MCMC with small values for *φ*_*X*_ and *φ*_*Y*_, post-processing of the MCMC samples often had no effect.

Estimates of all parameters from the small (with *L* = 500) and large datasets are summarized in table 1. In the small datasets, the introgression probabilities were *φ*_*X*_ ≈ 0.354 (with the CI 0.022–0.664) for the noncoding data and 0.280 (with CI 0.002–0.547) for the coding loci, while *φ*_*Y*_ was 0.104 (CI 0.000–0.306) for the noncoding data and 0.116 (CI 0.000–0.318) for the exonic data. When all loci from chromosome 1 were used, *φ*_*X*_ was 0.124 (with the CI 0.007–0.243) for the noncoding data and 0.161 (with CI 0.070– 0.264) for the exonic loci, while *φ*_*Y*_ was 0.048 (CI 0.000–0.139) for the noncoding data and 0.019 (CI 0.000–0.056) for the exonic data. The estimates were similar between the noncoding and exonic data, with greater proportions of migrants in *H. timareta* from *H. melpomene* than in the opposite direction. This was so despite the fact that *H. melpomene* had a smaller effective population size than *H. timareta*. Note that *H. melpomene* has a widespread geographical distribution whereas *H. timareta* is restricted to the Eastern Andes; the small *θ*_*M*_ estimates are most likely due to the fact that the *H. melpomene* sample was from a partially inbred strain to avoid difficulties with genome assembly. Estimates of *θ*s and *τ*s were smaller for the coding loci than for the noncoding loci, due to purifying selection removing deleterious nonsynonymous mutations.

Estimates of *φ*_*X*_ and *φ*_*Y*_ were different between the small and large datasets, but they involved large uncertainties, with the CIs for large datasets mostly inside the CIs for the small datasets. Another reason for the differences may be the variable rate of gene flow across the genome or chromosome. Note that *φ* in the MSci model reflects the long-term effects of gene flow and selection purging introgressed alleles, influenced by linkage to gene loci under natural selection (Martin and Jiggins, 2017). As a result, the introgression rates may be expected to vary across the genome.

### Analysis of data simulated under the double-BDI model of figure 6a

We conducted a small simulation to illustrate the feasibility of the double-BDI model (fig. 6), simulating 10 replicate datasets of *L* = 500, 2000, and 8000 loci. The three algorithms were used to process the MCMC samples, before they were summarized. A typical case is shown in figure 7 for the case of *L* = 500. While there are four unidentifiable towers in the 8-D posterior space (eq. 3), the MCMC visited only two of them, with different values for parameters around the BDI event at the node *ZW*. The dataset of *L* = 500 loci are very informative about the parameters for the BDI event at node *XY* (*φ*_*X*_, *φ*_*Y*_, *θ*_*X*_, *θ*_*Y*_), so that these had highly concentrated posteriors with well separated towers. We started the Markov chains with small values (e.g., 0.1 and 0.2) for *φ*_*X*_ and *φ*_*Y*_, so that the sampled points were all around the correct tower for those parameters. If the chain started with large *φ*_*X*_ and *φ*_*Y*_, it would visit a ‘mirror’ tower. Thus post-processing of the MCMC samples in the case of *L* = 500 mostly affected parameters around the BDI event at *ZW* (*φ*_*Z*_, *φ*_*W*_, *θ*_*Z*_, *θ*_*W*_). Figure 7 shows the effects on parameters *φ*_*Z*_ and *φ*_*W*_ using the *β*−*γ* algorithm. The CoG_0_ and CoG_*N*_ algorithms produced nearly identical results, and all algorithms were effective in removing label switching. The post-processed samples were summarized to calculate the posterior means and the HPD CIs (fig. 8).

**Figure 7:**
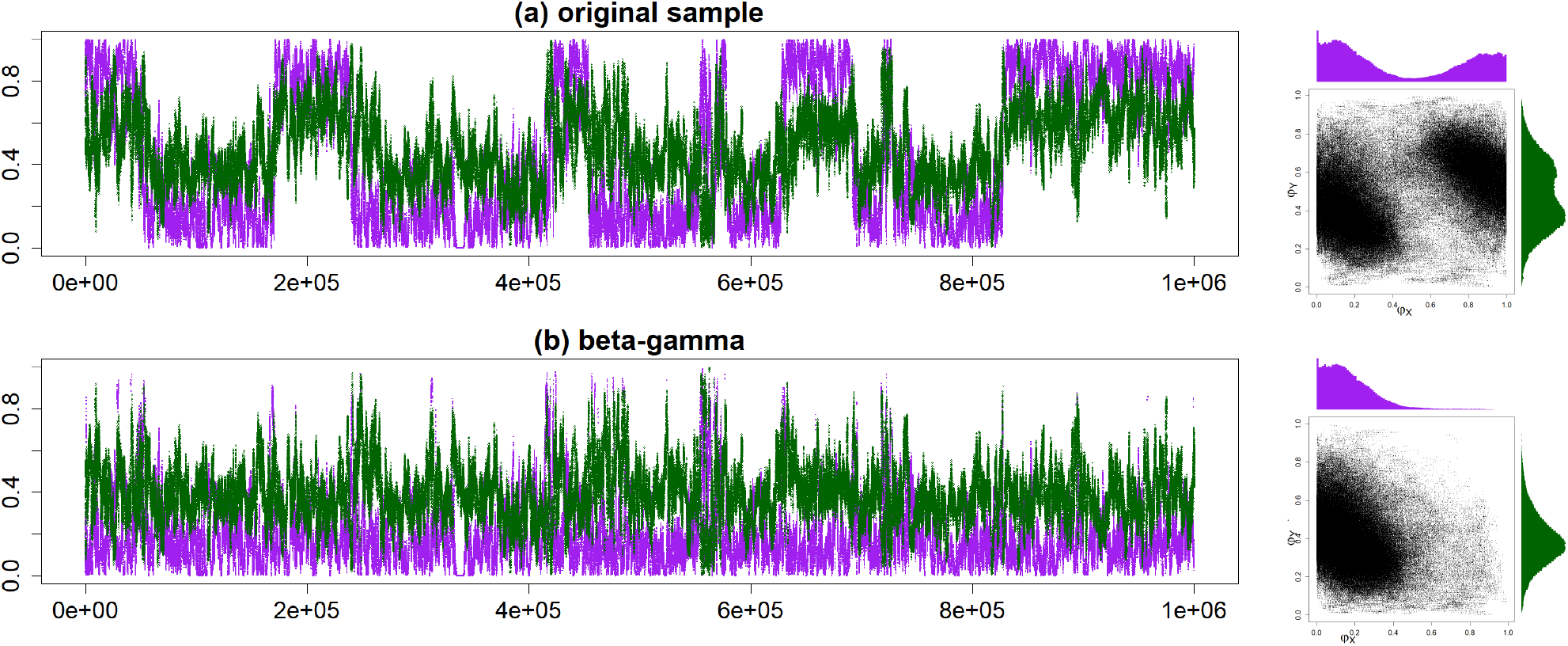
Trace plots of MCMC samples and 2-D scatter plots for parameters *φ*_*Z*_ (purple) and *φ*_*W*_ (green) (**a**) before and (**b**) after the post-processing of the MCMC samples for the double-DBI model of figure 6a. Post processing used the *β*−*γ* algorithm (**b**), while CoG_*N*_ and CoG_0_ produced nearly identical results (not shown). This is for replicate 2 for *L* = 500 loci (see fig. 8).

**Figure 8:**
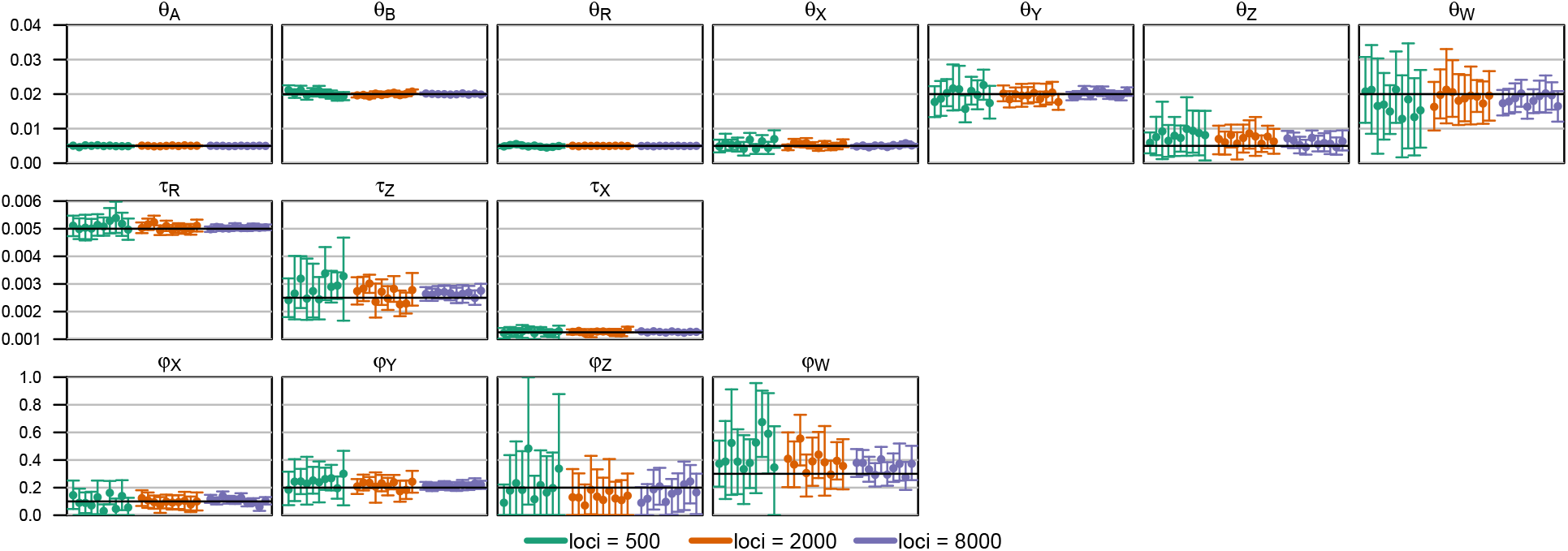
Posterior means and the 95% HPD CIs in 10 replicate datasets of *L* = 500, 2000, and 8000 loci, simulated and analyzed under the double-BDI model of figure 6a. The MCMC samples are post-processed using the *β*−*γ* algorithm before they are summarized (see fig. 7 for an example). Eight parameters are involved in the label-switching unidentifiability: *φ*_*X*_, *φ*_*Y*_, *θ*_*X*_, *θ*_*Y*_, *φ*_*Z*_, *φ*_*W*_, *θ*_*Z*_, and *θ*_*W*_ (see fig. 6).

At *L* = 2000 or 8000 loci, the four towers were well-separated and the MCMC visited only one of them. Applying the post-processing algorithms either had no effect or, in rare occasions, moved all sampled points from one tower to another.

Posterior means and the 95% highest-probability-density (HPD) credibility intervals (CI) for all parameters were summarized in figure 8. Parameters around the BDI event at *ZW* (*φ*_*Z*_, *φ*_*W*_, *θ*_*Z*_, *θ*_*W*_) are the most difficult to estimate. Nevertheless, the CIs for all parameters were smaller at *L* = 8000 than at *L* = 500 or 200, and the posterior means were converging to the true values. Note that while the simulation was conducted using one set of correct parameter values (say, Θ_1_ of fig. 6), we considered the estimates to be good if they were close to any of the four unidentifiable towers (say, Θ_2_, Θ_3_, or Θ_4_). This is analogous to treating the estimate as correct in Bayesian clustering if the true model includes two groups in proportions *p*_1_ = 10% and *p*_2_ = 90% with means *μ*_1_ = 100 and *μ*_2_ = 1, while the method of analysis infers two groups in proportions 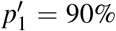 and 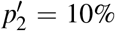 with means 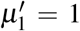 and 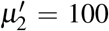. Just as Θ = (*p*_1_, *μ*_1_, *μ*_2_) and Θ′ = (*p*_2_, *μ*_2_, *μ*_1_) are unidentifiable towers and equally correct answers in the clustering problem, here Θ_1_, Θ_2_, Θ_3_, and Θ_4_ are equally correct answers.

### Analysis of data simulated with one BDI event with poorly separated modes

We simulated a more challenging dataset for the relabelling algorithms, with *L* = 500 loci, under the BDI model of figure 1a with (*φ*_*X*_, *φ*_*Y*_) = (0.7, 0.2) (see table S1). As *φ*_*X*_ and *φ*_*Y*_ were not too far away from 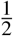 and the dataset is small, the posterior modes were poorly separated, with considerable mass near 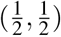. In the unprocessed MCMC sample, *φ*_*X*_ had two modes around 0.8 and 0.2 and the chain was switching between them (fig. S7a). The posterior means were at 0.51 for *φ*_*X*_ and 0.50 for *φ*_*Y*_, close to 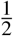 (fig. S7a). These were misleading summaries, as the sample was affected by label switching. In the processed samples (fig. S7b-d), label switching was successfully removed and both *φ*_*X*_ and *φ*_*Y*_ were single-moded. The three algorithms (*β*−*γ*, CoG_*N*_, and CoG_0_) produced similar results, with single-moded posterior, around the tower (*φ*_*X*_, *φ*_*Y*_) = (0.7, 0.2). The posterior means of (*φ*_*X*_, *φ*_*Y*_) were (0.755, 0.447), (0.766, 0.461), and (0.765, 0.462) for the three algorithms, *β*−*γ*, CoG_*N*_, and CoG_0_, respectively (table S1). The estimates from *β*−*γ* were slightly closer to the true values than those from CoG_*N*_ and CoG_0_. The three relabelling algorithms worked well even when the posterior modes were poorly separated.

The parameters in the model not involved in the label-switching, such as the species divergence and introgression times (*τ*_*R*_, *τ*_*X*_) and the population sizes for the modern species and for the root (*θ*_*A*_, *θ*_*B*_, *θ*_*R*_), were well estimated, with the posterior means close to the true values and with narrow CIs (table S1). However, the parameters involved in the label switching (*φ*_*X*_, *φ*_*Y*_, *θ*_*X*_, *θ*_*Y*_) were poorly estimated at this data size (with *L* = 500 loci), because of the difficulty to separate the two towers and the influence of the priors. The estimates should improve if more loci are used in the data. To confirm this expectation, we simulated two more datasets with *L* = 2000 and 8000 loci, respectively. In those two larger datasets, parameters not involved in label switching (*τ*_*R*_, *τ*_*X*_, *θ*_*A*_, *θ*_*B*_, *θ*_*R*_) had very narrow CIs (table S1). The posterior means of Θ = (*φ*_*X*_, *φ*_*Y*_) were closer to the true values (0.7, 0.2), and the 95% CIs were narrower than in the small dataset of *L* = 500 (table S1). Note that ancestral population sizes (such as *θ*_*X*_ and *θ*_*Y*_) are hard to estimate even in models of unidirectional introgression which do not have label-switching issues (Huang *et al*., 2020).

## DISCUSSION

### Data size, precision of parameter estimation, MCMC convergence, and the utility of the relabelling algorithms

We have observed three kinds of behaviors of the MCMC algorithm and the relabelling algorithms depending on the data size. In small datasets, the parameters are poorly estimated with large uncertainties, and the posterior modes (the unidentifiable towers) are not well separated. In such cases, applying simple constraints (such as 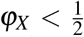) is problematic because the truncation distorts the marginal summaries of the posterior, with different constraints producing different posterior summaries (Richardson and Green, 1997; Celeux *et al*., 2000; Stephens, 2000). The relabelling algorithms are preferable. An example is the small dataset of *L* = 500 loci simulated under the model of one BDI event (fig. S7, table S1).

In intermediate datasets, the parameters are well estimated, the posterior modes are well separated, but the MCMC algorithm jumps between the modes, switching labels. In such cases, any of the relabelling algorithms will work well. If the posterior modes are far away from the boundary defined by the constraints (such as 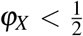), even simple constraints will work well. Examples include two small butterfly datasets (figs. 3 & S1), and the datasets simulated under the double BDI model (fig. 7).

Finally, in very large datasets, the parameters are extremely well estimated with very narrow CIs, and the posterior modes are so sharply concentrated that the MCMC algorithm stays on one of the unidentifiable towers and never moves to the mirror towers. Furthermore, in multiple runs of the same analysis the MCMC may be ‘stuck’ on different towers. In such cases, the relabelling algorithms will either not move any sample points at all or move all points from one tower to another, and any of the algorithms will work well. This scenario is common in analyses of large genomic datasets with thousands of loci, such as the large noncoding and exonic datasets from the *Heliconius* butterflies (fig. 2); See Finger *et al*. (2021) and Thawornwattana *et al*. (2021) for many more examples.

We note that in all three scenarios, the relabelling algorithms (in particular, the *β*−*γ* algorithm) were either better or not worse than the alternatives such as imposing simple constraints. Given that even the *β*−*γ* algorithm involves minimal computation, we recommend its automatic use in all cases. Samples from different runs visiting different unidentifiable modes may be merged before post-processing using the relabelling algorithm.

In theory, if the MCMC has converged and is mixing well and the algorithm is run long enough, it should visit the unidentifiable towers with exactly the same probability and the means of introgression probabilities from the unprocessed samples should be 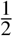. One might expect this expectation to provide a useful criterion for diagnosing the convergence of such MCMC algorithms. Indeed Jasra *et al*. (2005) regarded it “a *minimum* requirement of convergence for a mixture posterior to be such that we have explored all possible labellings of the parameters”. Here the labellings correspond to the unidentifiable towers. We suggest that this requirement is too stringent and unnecessary. As discussed above, in large genomic datasets, the posterior may be highly concentrated, and the chain may never jump between the towers even in very long MCMC runs. While the chain may be visiting different mirror towers in different runs of the same analysis, each chain may be exploring the space around the visited tower thoroughly, and after label switching is removed, the MCMC samples from the different runs may produce nearly identical posterior summaries, suggesting that reliable inference is entirely possible. In simulations of large datasets, the posterior estimates after label switching problems are removed converge to the true values (e.g., Flouri *et al*., 2020, fig. S10A). We suggest that exploration of all unidentifiable towers is unnecessary for correct inference and should not be used as a criterion for diagnosing MCMC convergence. Instead convergence diagnosis should be applied after the MCMC sample is processed to remove label switching. For example, one should run the same analysis multiple times and confirm that the posterior summaries when the MCMC samples are processed and mapped onto the same tower are consistent between runs. The efficiency of the MCMC algorithm or the effective sample size (ESS) (Yang and Rodríguez, 2013) should also be calculated using the processed samples.

### Identifiability of MSci models with unidirectional introgressions

The identifiability of MSci models involving unidirectional introgression (UDI) events appears to be simpler than for BDI models (Flouri *et al*., 2020; Jiao *et al*., 2021). MSci model A (Flouri *et al*., 2020) is consistent with three different biological scenarios (fig. 9a-c). In scenario A_1_, two species *SH* and *TH* merge to form a hybrid species *HC*, but the two parental species become extinct after the merge. This scenario may be rare. In scenario A_2_, species *SUX* contributes migrants to species *THC* at time *τ*_*H*_ and has since become extinct or is unsampled in the data. In scenario A_3_, *TUX* is the extinct or unsampled ghost species. The three scenarios are unidentifiable using genomic data. Model B_1_ assumes introgression from species *RA* to *TC* at time *τ*_*S*_ = *τ*_*H*_ (fig. 9d). This is distinguishable using genetic data from the alternative model B_2_ in which there is introgression from *RB* to *SC* (fig. 9e). Note that models B_1_ and B_2_ are both special cases of model A_1_ with different constraints (that is, *τ*_*S*_ = *τ*_*H*_ *< τ*_*T*_ for model B_1_ and *τ*_*S*_ *> τ*_*H*_ = *τ*_*T*_ for model B_2_).

**Figure 9:**
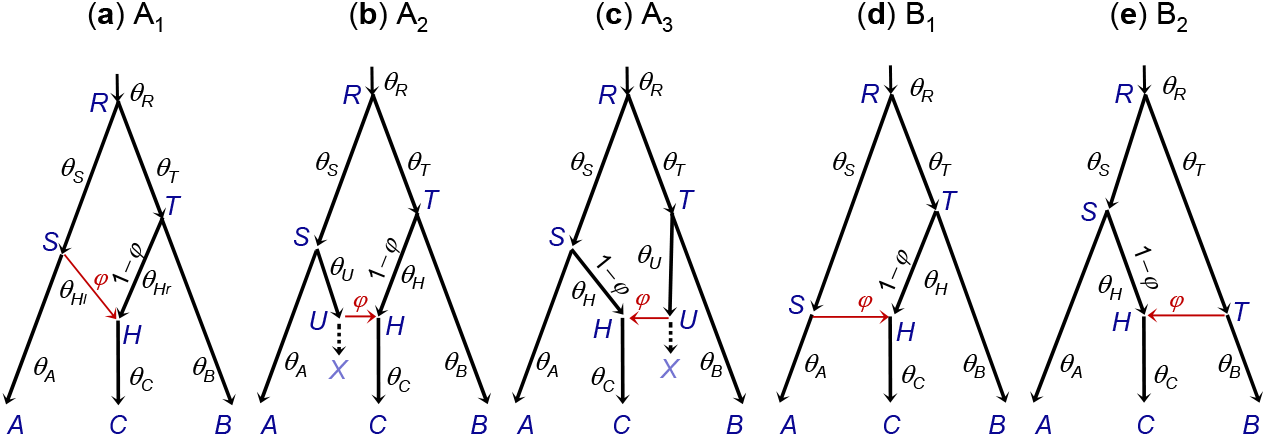
Species trees for three species (*A, B*, and *C*) illustrating MSci models of types A and B of Flouri *et al*. (2020, fig. 1). (**a**-**c**)Three interpretations of MSci model A are indistinguishable/unidentifiable. (**d, e**) Two versions of MSci model B are identifiable.

Note that the sampling configuration may affect the identifiability of parameters in the model (Yu *et al*., 2012; Zhu and Degnan, 2017). The simplest such example may be the population size parameter (*θ*). If at most one sequence per locus is sampled from a species, the population size for that species will be unidentifiable. Similarly, if no more than one sequence per locus can enter an ancestral population when we trace the genealogy of the sampled sequences backwards in time, *θ* for that ancestral species will be unidentifiable. Such unidentifiability disappears when multiple sequences per species are sampled. Note that a diploid sequence is equivalent to two haploid sequences. Similarly introgression models that are unidentifiable with one sampled sequence per species may become identifiable when multiple sequences per species are sampled (Zhu and Degnan, 2017).

Even if the model is mathematically identifiable with one sequence per species per locus, including multiple samples per species (in particular, species that are descendants of a hybridization node in the species tree) can boost the information content in the data dramatically. Thus we recommend the use of multiple samples per species in studies of cross-species gene flow, and suggest that the most interesting scenario for studying unidentifiability of models of gene flow should be full likelihood analysis of multilocus sequence data, with multiple sequences sampled per species.

It is noteworthy that many parameter settings and data configurations exist in which some parameters are hard to estimate, because the data lack information about them. For example, ancestral population sizes for short and deep branches in the species tree are hard to estimate, because most sequences sampled from modern species may have coalesced before reaching that population when we trace the genealogy of the sample backwards in time (Huang *et al*., 2020). Similarly, if few sequences reach a hybridization node, there will be little information in the data about the introgression probabilities at that node. In such cases, even if the model is identifiable mathematically, it may be nearly impossible to estimate the parameters with any precision even with large datasets.

In some cases, certain parameters may be nearly at the boundary of the parameter space, and this may create near unidentifiability with multiple modes in the posterior. For example, two speciation events that occur in rapid succession will generate a very short branch in the species tree with a near trichotomy in the species tree. Then MSci models that posit the same introgression events but different histories of species divergences will fit the data nearly equally well and become multiple modes in the posterior space (see Finger *et al*., 2021 for an example). Similarly introgression probabilities near 0 or 1 can also create nearly equally good explanations of the data and become multiple modes in the posterior. In such situations, the MCMC samples around different modes should be summarized separately.

### Unidentifiability of heuristic methods

As mentioned in Introdution, the unidentifiability discussed in this paper concerns the intrinsic nature of the inference problem when introgression models are applied to genomic sequence data, and thus applies to not only full likelihood methods but also heuristic methods based on summaries of the sequence data. Indeed a model that is unidentifiable by a full likelihood method must be unidentifiable by any heuristic method. In contrast, a model that is identifiable by a full likelihood method may still be unidentifiable by a heuristic method as the heuristic method may not be using all information in the data. Here we briefly discuss a few heuristic methods, focusing on their common features. Interested readers may consult the recent reviews by Elworth *et al*. (2019) and Hibbins and Hahn (2021). Heuristic methods developed up to now are mostly of two kinds, based on either genome-wide averages or estimated gene trees for genomic segments (loci).

The popular *ABBA*-*BABA* test (Durand *et al*., 2011) uses the parsimony-informative site patterns across the genome to detect gene flow. Consider three populations/species *S*_1_, *S*_2_, and *S*_3_, with the given phylogeny ((*S*_1_, *S*_2_), *S*_3_), plus an outgroup species *O*. There are three parsimony-informative site patterns: *BBAA, ABBA*, and *BABA*. Here *A* and *B* represent any two distinct nucleotides and *BBAA* means *S*_1_ and *S*_2_ have the same nucleotide while *S*_3_ and *O* have another. For very closely related species, one may consider nucleotide *A* in the outgroup as the ancestral allele and *B* the derived allele. Site pattern *BBAA* matches the species tree, while *ABBA* and *BABA* are the mismatching patterns. Given the species tree with no gene flow, the two mismatching patterns have the same probability, but when there is gene flow between *S*_1_ (or *S*_2_) and *S*_3_, they will have different probabilities. The difference between the two mismatching site-pattern counts can then be used to test for the presence of gene flow (Durand *et al*., 2011):

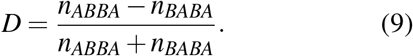

The *D*-statistic may also be seen as a comparison between the number of derived alleles shared by *S*_2_ and *S*_3_ with that shared by *S*_1_ and *S*_3_. The test has more power to detect inflow (gene flow from *S*_3_ → *S*_2_) than outflow (from *S*_2_ → *S*_3_) (Hibbins and Hahn, 2021, fig. S3). It can test for the presence of gene flow, but provides no information about its direction, timing or strength.

A number of variants or extensions of the *D*-statistic have been proposed. Instead of the parsimony-informative site patterns, the average sequence distance between species may be used to construct a similar test (Hahn and Hibbins, 2019). The site pattern counts can also be used to estimate the introgression probability, as in the program HYDE (Blischak *et al*., 2018; Kubatko and Chifman, 2019):

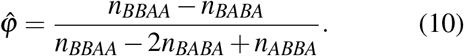

This is based on the hybrid speciation model (with *τ*_*S*_ = *τ*_*H*_ = *τ*_*T*_ and *θ*_*S*_ = *θ*_*T*_ in model A_1_ of fig. 9). The estimate may be biased if this symmetry assumption does not hold.

The *D*-statistic has been extended to the case of five species, with a symmetric species tree assumed, in the so-called *D*_FOIL_ test, with the aim to detect the direction of gene flow (Pease and Hahn, 2015). Another extension is by Hamlin *et al*. (2020), to include the site pattern *BBAA* to form

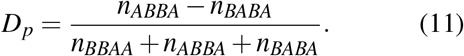

Interpreted as an estimate of the genome-wide introgression proportion (*φ*), *D*_*p*_ has a negative bias, which is more serious for outflow (from *S*_2_ → *S*_3_) than for inflow (with gene flow from *S*_3_ → *S*_2_) (Hamlin *et al*., 2020, fig. 3).

Note that both the site-pattern counts and between-species distances are genome-wide averages. If the data consist of multi-locus sequence alignments, they can be merged (concatenated) into a super-alignment to calculate those statistics. A great advantage of those methods is that they involve minimal computation. A serious drawback is that they do not make use of information in genealogical variations across the genome. Like the coalescent process, gene flow between species creates stochastic fluctuations in the genealogical history (gene tree topology and coalescent times) across the genome, with the probability distribution given by the parameters in the multispecies coalescent model with gene flow, including species divergence times, effective population sizes for modern and ancestral species, and the directions and rates of gene flow. As a result, there is important information about those parameters in such genomic variation, but this information is ignored by those methods. Under the assumption of strong linkage among sites in the same genomic segment (locus), all sites at the same locus share the genealogical history while differences among sites of the same locus reflect the stochastic fluctuations of the mutation process. In calculations of genome-wide averages, those two sources of variation are confounded, and the information on coalescent fluctuations among loci is lost (Shi and Yang, 2018; Zhu and Yang, 2021). As a result, most parameters in the coalescent model with introgression are unidentifiable by the heuristic methods mentioned above. None of them can detect signals of gene flow between sister species, and for non-sister species, none of them can estimate the introgression probabilities when gene flow occurs in both directions (e.g., *φ*_*X*_ and *φ*_*Y*_ in fig. 1a or *α* and *β* in fig. S3a).

The second kind of heuristic methods use reconstructed gene tree topologies for multiple loci as the input data. Consider again the species quartet *S*_1_, *S*_2_, *S*_3_, and *O* (outgroup), with the given phylogeny ((*S*_1_, *S*_2_), *S*_3_), and one sampled sequence per species. The two mismatching gene trees ((*S*_2_, *S*_3_), *S*_1_) and ((*S*_3_, *S*_1_), *S*_2_) have the same probability if there is coalescence but no gene flow, but different probabilities if there is in addition gene flow between the non-sister species (between *S*_1_ and *S*_3_ or between *S*_2_ and *S*_3_). Thus the frequencies of gene tree topologies can be used to estimate the introgression probability, as in the SNAQ method (Solis-Lemus and Ane, 2016, see also Yu *et al*., 2012). As there are only two free quantities (frequencies of three gene trees with the sum to be 1), the approach can estimate the internal branch length in coalescent units and the introgression probability, but not any other parameters in the model.

In the general case, the probabilities of gene tree topologies under any introgression model can be calculated by summing over the compatible coalescent histories (Yu *et al*., 2012, 2014). The probability distribution of gene tree topologies can then be used to distinguish among different introgression models and to estimate the parameters in the introgression model by ML (as in PhyloNet; Cao *et al*., 2019), treating gene tree topologies as the data. A concern with the two-step method is that the estimated gene trees may involve uncertainties or errors, in particular when the species are closely related. Similarly, including genetree branch lengths (coalescent times) makes many introgression models that are unidentifiable based on gene tree topologies alone become identifiable (Yu *et al*., 2012; Zhu and Degnan, 2017). However, two step methods that make use of estimated branch lengths do not have good performance as the large uncertainties and errors in the estimated branch lengths can have a major impact on inference of species divergence and cross-species gene flow (Degnan, 2018).

Pardi and Scornavacca (2015) studied the unidentifiability of network models using data of gene tree topologies ‘displayed’ by the network (fig. 10). Binary species trees generated by taking different parental paths at hybridization nodes are called “displayed species trees” (Pardi and Scornavacca, 2015) or “parental species trees” (Kubatko, 2009). For example, the two network models *N*_1_ and *N*_2_ of figure 10a are unidentifiable because they induce the same three displayed species trees with the same branch lengths (Pardi and Scornavacca, 2015). However, as pointed out by Zhu and Degnan (2017), *N*_1_ and *N*_2_ are identifiable using gene tree topologies if multiple sequences are sampled from *B*.

**Figure 10:**
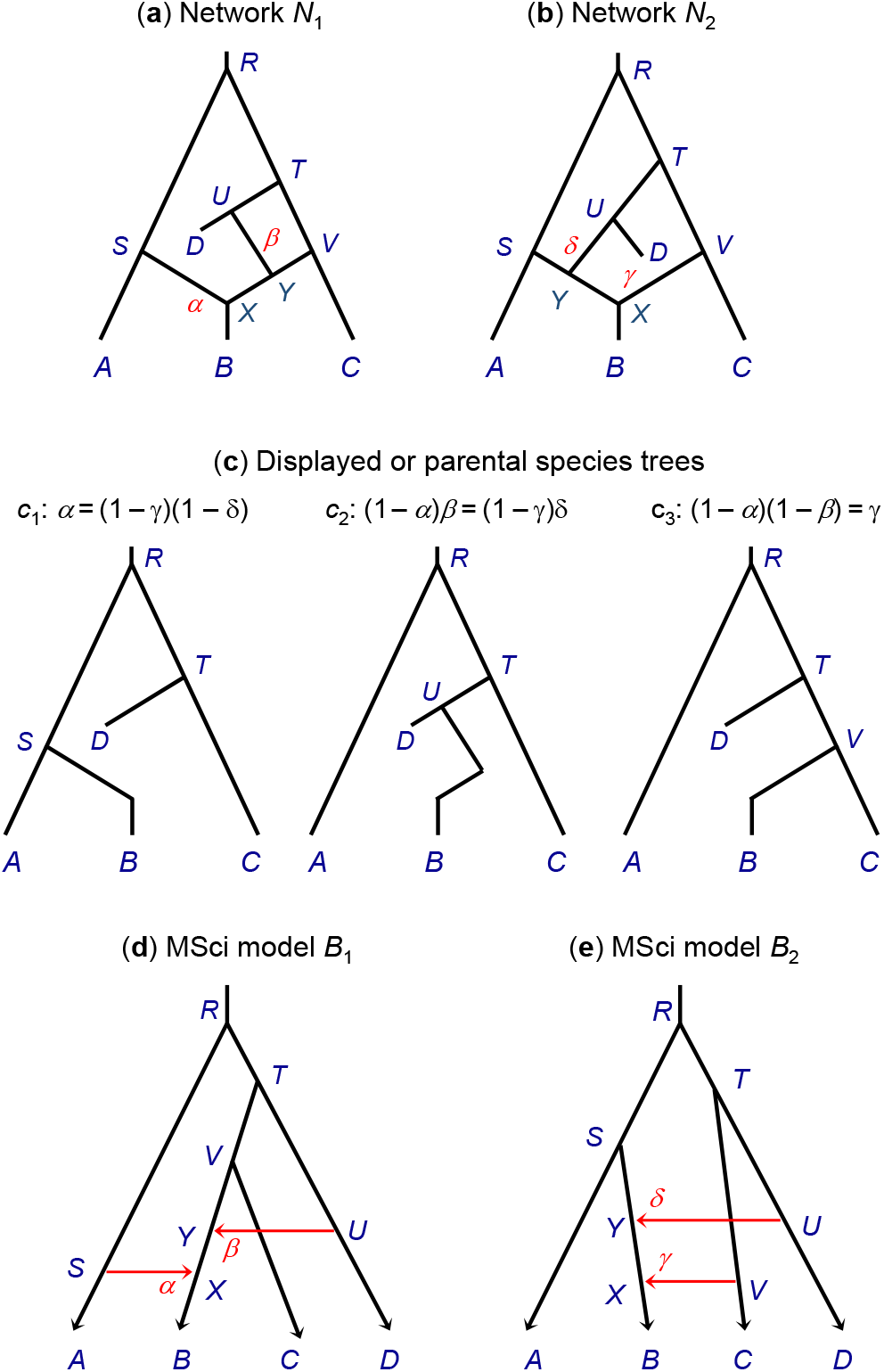
(**a&b**) Two phylogenetic networks for four species (*A, B,C, D*), each with two hybridization events from Pardi and Scornavacca (2015) that are unidentifiable using gene tree topologies with one sequence sampled per species. (**c**) Network *N*_1_ gives rise to three ‘displayed species trees’ in probabilities *α*, (1 − *α*)*β*, and (1 − *α*)(1 − *β*), while *N*_2_ gives rise to the same three displayed species trees with probabilities (1 − *γ*)(1 − *δ*), (1 − *γ*)*δ*, and *γ*. The two networks thus give the same distribution of gene tree topologies, and are thus unidentifiable. However, *N*_1_ and *N*_2_ are identifiable when multiple samples are taken from species *B*. (**d&e**) MSci models corresponding to networks *N*_1_ and *N*_2_. With information from branch lengths (coalescent times) and using multilocus sequence data, those models are identifiable by full likelihood method, as are the 18 parameters in each model, including five species divergence/introgression times (*τ*s), eleven population sizes (*θ*s), and two introgression probabilities.

Previously (Kubatko, 2009, eq. 3) formulated the probability distribution of gene trees (topology alone or topology with coalescent times) as a mixture over the displayed species trees. To simulate gene trees or sequence data at a locus, one samples a displayed species tree first and then simulates the gene tree and sequence alignment according to the simple MSC model (Gerard *et al*., 2011). This formulation is correct only in the special case where each hybridization node on the species tree has at most one sequence from all its descendant populations (Zhu and Degnan, 2017). In general it is incorrect, as it forces all sequences at the locus to take the same parental path at each hybridization node, whereas correctly there should be a binomial sampling process when two or more sequences reach a hybridization node. In model *N*_1_ of figure 10a, when multiple *B* sequences reach species *X*, it should be possible for some sequences to take the left parental path while the others take the right path.

Even though the notion that gene trees are displayed by a phylogenetic network has played a central role in many studies that attempt to use gene tree topologies to construct the phylogenetic network, examination of the displayed gene trees is not a reliable approach to studying the unidentifiability of phylogenetic network models (Zhu and Degnan, 2017). The most probable gene tree may even have a topology that is different from all of the displayed trees (Zhu and Degnan, 2017). As suggested by Zhu and Degnan (2017), one should instead explicitly treat the biological process of coalescent and introgression in the model. We also suggest that multiple sequences be sampled per species (in particular from species involved in hybridization or from descendant species of hybridization nodes) when genomic data are used to infer gene flow. Note that both MSci models corresponding to networks *N*_1_ and *N*_2_ are identifiable when genomic sequence data with multiple samples per species are analyzed using full likelihood methods (fig. 10d&e), as are all parameters in each models (fig. 10a′&b′).

We note that there is currently a wide gap between likelihood and heuristic methods. Heuristic methods are computationally orders-of-magnitude faster than likelihood methods, which in particular do not scale well for large genomic datasets. The statistical properties of heuristic methods are also incomparably poorer than those of likelihood methods: heuristic methods are simply unable to provide any estimates for many fundamental population parameters for characterizing the evolutionary history of the species, such as the species divergence/introgression times, the population sizes of extant and extinct species, the introgression probabilities, etc. There is an acute need for improving the statistical performance of the heuristic methods and the computational efficiency of the full likelihood methods.

Given the limitations of the heuristic methods, one should apply caution when using them to draw biological conclusions concerning gene flow between species. For example, does gene flow occur more often between sister species or between non-sister species? When gene flow occurs between two species, does it often involve one direction (UDI) or both directions (BDI)? Most heuristic methods cannot identify or detect gene flow between sister species or gene flow in both directions, but it may be erroneous to conclude that such gene flow never occurs in nature. Whether BDI or UDI is more common is an interesting empirical question, but both models provide important biological hypotheses testable using genomic sequence data. In a recent analysis of genomic sequence data from the North American chipmunks (*Tamias quadrivittatus*), the use of the *D*-statistic and HyDe detected no evidence of gene flow affecting the nuclear genome despite widespread mitochondrial gene flow (Sarver *et al*., 2021). However a reanalysis of the same data using BPP revealed robust evidence for multiple ancient introgression events, involving both sister and nonsister species (Ji *et al*., 2021).

### Estimation of introgression probabilities despite unidentifiability

The three algorithms for post-processing MCMC samples under the BDI model produced very similar results in the applications in this study. In particular the simple center-of-gravity algorithms produced results that appear to be as good as the more elaborate *β*−*γ* algorithm, despite the fact that the normal distribution is a poor approximation to the posterior of introgression probabilities (*φ*_*X*_ and *φ*_*Y*_). This is due to the fact that the distributions (or the distances in the CoG algorithms) are used to compare the unidentifiable mirror positions of sample points only, but are not used to approximate the posterior distribution of those parameters, which are estimated by using the processed samples. For the same reasons, if there exist multiple modes in the posterior that are not due to label switching, such genuine multimodality will not be removed by the relabelling algorithms (Stephens, 2000). Similarly, while we fit independent distributions for parameters in the algorithms (eq. 6), there is no need to assume independence in the posterior for the algorithms to work.

A model with a label-switching type of unidentifiability can still be applied in real data analysis. In the clustering problem, the Bayesian analysis may reveal the existence of two groups, in proportions *p*_1_ and *p*_2_ = 1 − *p*_1_ with means *μ*_1_ and *μ*_2_, and it does not matter if it cannot decide which group should be called ‘group 1’. The twin towers Θ and Θ′ under the BDI model of figure 1 constitute a mathematically similar label-switching problem. However, Θ and Θ′ may represent different biological scenarios or hypotheses. Suppose that species *A* and *B* are distributed in different habitats (dry for *A* and wet for *B*, say), and suppose the ecological conditions have changed little throughout the history of the species. Θ with 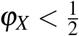 and 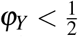 may mean that species *A* has been in the dry habitat over the whole time period since species divergence at time *τ*_*R*_, while species *B* has been in the wet habitat, and they came into contact and exchanged migrants at time *τ*_*X*_. In contrast, Θ′ with 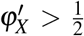 and 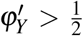 may mean that species *A* was in the wet habitat and species *B* was in the dry habitat since species divergence at time *τ*_*R*_, but when they came into contact at time *τ*_*X*_ they nearly replaced each other, switching places, so that today species *A* is found in the dry habitat while *B* in the wet habitat. The two sets of parameters Θ and Θ′ may thus mean different biological hypotheses. As genomic data from modern species provide information about the order and timings of species divergences and cross-species introgressions, but not about the geographical locations and ecological conditions in which the divergences and introgressions occurred, such biological scenarios are unidentifiable using genomic data and become unidentifiable towers in the posterior distribution in Bayesian analysis of genomic data under the MSci model. Unidentifiable models discussed in this paper are all of this nature. The algorithms we developed in this paper remove label switching in the MCMC sample, but do not remove the unidentifiability of the BDI models. The researcher has to be aware of the unidentifiability and use external information to choose between such equally supported explanations of the genomic data.

In the above example, the scenario of near-complete replacement represented by Θ′ may be implausible for most systems, and in our relabelling algorithms, we start with small *φ*_*X*_ and *φ*_*Y*_ as much as possible (through the initial condition *φ*_*X*_ + *φ*_*Y*_ *<* 1). When the introgression probabilities are intermediate, the biological interpretations may not be so clear-cut, but unidentifiability exists nevertheless. In the example of figure S7 and table S1 for the simulated data with one BDI event, the choice between the two unidentifiable towers Θ = (*φ*_*X*_, *φ*_*Y*_) = (0.7, 0.2) and Θ′ = (0.3, 0.8) may not be easy.

In the current implementation of BDI models in BPP, each branch in the species tree is assigned its own population size parameter (Flouri *et al*., 2020). We note that if all species on the species tree are assumed to have the same population size (*θ*), unidentifiability persists. However, if we assume that the population size remains unchanged by the introgression event: e.g., *θ*_*X*_ = *θ*_*A*_ and *θ*_*Y*_ = *θ*_*B*_ in figure 1, the model becomes identifiable. The assumption of the same population size before and after an introgression event appears to be plausible biologically. It reduces the number of parameters by two for each BDI event, and removes unidentifiability. It may be worthwhile to implement such models. At any rate, the relabelling algorithms we have implemented makes it possible to apply the BDI models to genomic sequence data despite their unidentifiability.

## METHODS AND MATERIALS

### Introgression in *Heliconius* butterflies

We fitted the BDI model to the genomic sequence data for three species of *Heliconius* butterflies: *H. melpomene, H. timareta*, and *H. numata* (Consortium, 2012; Martin *et al*., 2013). The species tree or MSci model assumed is shown in figure 2, with introgression between *H. melpomene* and *H. timareta*. The two species are known to hybridize, although no attempt has yet been made to infer the direction or strength of introgression (except for colour-pattern genes; Martin *et al*., 2013). There are 31,166 autosomal noncoding loci and 36,138 autosomal exonic loci, with one diploid sequence sampled per species per locus. The sequence length ranges from 11 to 991 bps (median 93) for the noncoding loci and from 11 to 10,672 bps (median 112) for the exonic loci. We used chromosome 1, which has 2592 noncoding and 3023 exonic loci, and analyzed either the first 500 loci or all the loci on the chromosome. Note that a diploid sequence from each species is equivalent to two haploid sequences, so that the population size parameter (*θ*) for that species is estimable. Heterozygotes in the diploid sequence are represented by IUPAC ambiguity codes (e.g., with Y meaning a T/C heterozygote) and resolved into compatible nucleotides in BPP using an analytical integration algorithm (Gronau *et al*., 2011; Yang, 2015; Flouri *et al*., 2018), which averages over all possible genotypic phase resolutions of heterozygote sites, weighting them according to their likelihood based on the sequence alignment at the locus. In simulations, this approach had indistinguishable performance from analyzing fully phased genomic sequences (Gronau *et al*., 2011; Huang *et al*., 2021).

We used gamma priors for the population sizes (*θ*) and for the age of the root (*τ*_0_): *θ* ∼ *G*(2, 400) with the mean 0.005 substitution per site, and *τ* ∼ *G*(2, 400) with mean 0.005. The introgression probabilities were assigned beta priors *φ*_*X*_, *φ*_*Y*_ ∼ *B*(1, 1), which is the uniform 𝕌(0, 1). We used a burn-in of 16000 iterations, and then took 2 × 10^5^ samples, sampling every 5 iterations. Running time on a server using 9 threads of Intel Xeon Gold 6154 CPU (3.0GHz) was about 1 hour for the small datasets and 10 hours for the large ones.

Convergence of the MCMC algorithms was assessed by checking for consistency between independent runs, taking into account possible label-switching issues.

### Simulation under the double-BDI model

We simulated and analyzed data to under the double-BDI model of figure 6. Gene trees with branch lengths (coalescent times) were simulated under the MSci model and given the gene trees, sequences were evolved along the branches on the gene tree under the JC model (Jukes and Cantor, 1969). The parameters used were *φ*_*X*_ = 0.1, *φ*_*Y*_ = 0.2, *φ*_*Z*_ = 0.2, *φ*_*W*_ = 0.3, *τ*_*R*_ = 0.005, *τ*_*Z*_ = *τ*_*W*_ = 0.0025, *τ*_*X*_ = *τ*_*Y*_ = 0.00125, *θ*_*R*_ = *θ*_*Z*_ = *θ*_*X*_ = *θ*_*A*_ = 0.005, and *θ*_*W*_ = *θ*_*Y*_ = *θ*_*B*_ = 0.02. Each dataset consisted of *L* = 500, 2000 and 8000 loci, with *S* = 16 sequences per species per locus, and with the sequence length to be 500 sites. The number of replicate datasets was 10.

The data were then analyzed using BPP under the double-BDI model (fig. 6) to estimate the 14 parameters. We use gamma priors *τ*_0_ ∼ *G*(2, 400) for the root age with the mean to be the true value (0.005), and *θ* ∼ *G*(2, 200) with the mean 0.01 (true values are 0.005 and 0.02). We used a burn-in of 32,000 iterations, and then took 5 × 10^5^ samples, sampling every 2 iterations. Analysis of each dataset took ∼ 10hrs for *L* = 500 and ∼ 130hrs for *L* = 8000, using 8 threads on a server. The MCMC samples were processed to remove label-switching problems before they were summarized to approximate the posterior distribution.

### Simulation under a BDI model with poorly separated towers

We simulated a small dataset, with *L* = 500 loci, under the BDI model of figure 1a, with (*φ*_*X*_, *φ*_*Y*_) = (0.7, 0.2) (see table S1 for the true values of all parameters). As *φ*_*X*_ and *φ*_*Y*_ were not far away from 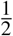 and the dataset was small, the posterior of the parameters was expected to be diffuse, and the posterior modes for parameters involved in the label-switching (or the two unidentifiable towers) to be poorly separated, posing a challenge to our relabelling algorithms.

We assigned gamma priors *τ*_0_ ∼ *G*(2, 200) for the root age with the mean to be the true value (0.01), and *θ* ∼ *G*(2, 400) with the mean 0.005 (true values are 0.002 and 0.01). We used a burn-in of 32,000 iterations, and then took 2 × 10^5^ samples, sampling every 10 iterations. We run the same analysis twice to confirm consistency between runs, after the MCMC samples were processed to remove label switching.

## Acknowledgements

James Mallet and Fernando Seixas provided the genomic datasets for the *Heliconius* butterflies. We thank James Mallet, Yuttapong Thawornwattana, and Christopher Blair for comments on an earlier version of the paper, and Xiyun Jiao for discussions. This study has been supported by Biotechnology and Biological Sciences Research Council grant (BB/T003502/1) and a BBSRC equipment grant (BB/R01356X/1).

## ONLINE SUPPORTING INFORMATION (SI)

- Figure S1: Analysis of the first 500 exonic loci of the *Heliconius* data.
- Figure S2: Three models with a BDI event between sister species.
- Figure S3: Two models with a BDI event between nonsister species.
- Figure S4: Three models with a BDI event between nonsister species.
- Figure S5: Two BDI events between non-sister species creating four unidentifiable models.
- Figure S6: Scatterplots illustrating the CoGN_0_ algorithm.
- Figure S7: Tracecatter plots for *φ*_*X*_ and *φ*_*Y*_ in analysis of a dataset of *L* = 500 loci simulated under the BDI model of figure 1.
- Table S1: Posterior means and 95% HPD CIs for parameters in the BDI model from a simulated data of *L* = 500 loci.

**Figure S1:**
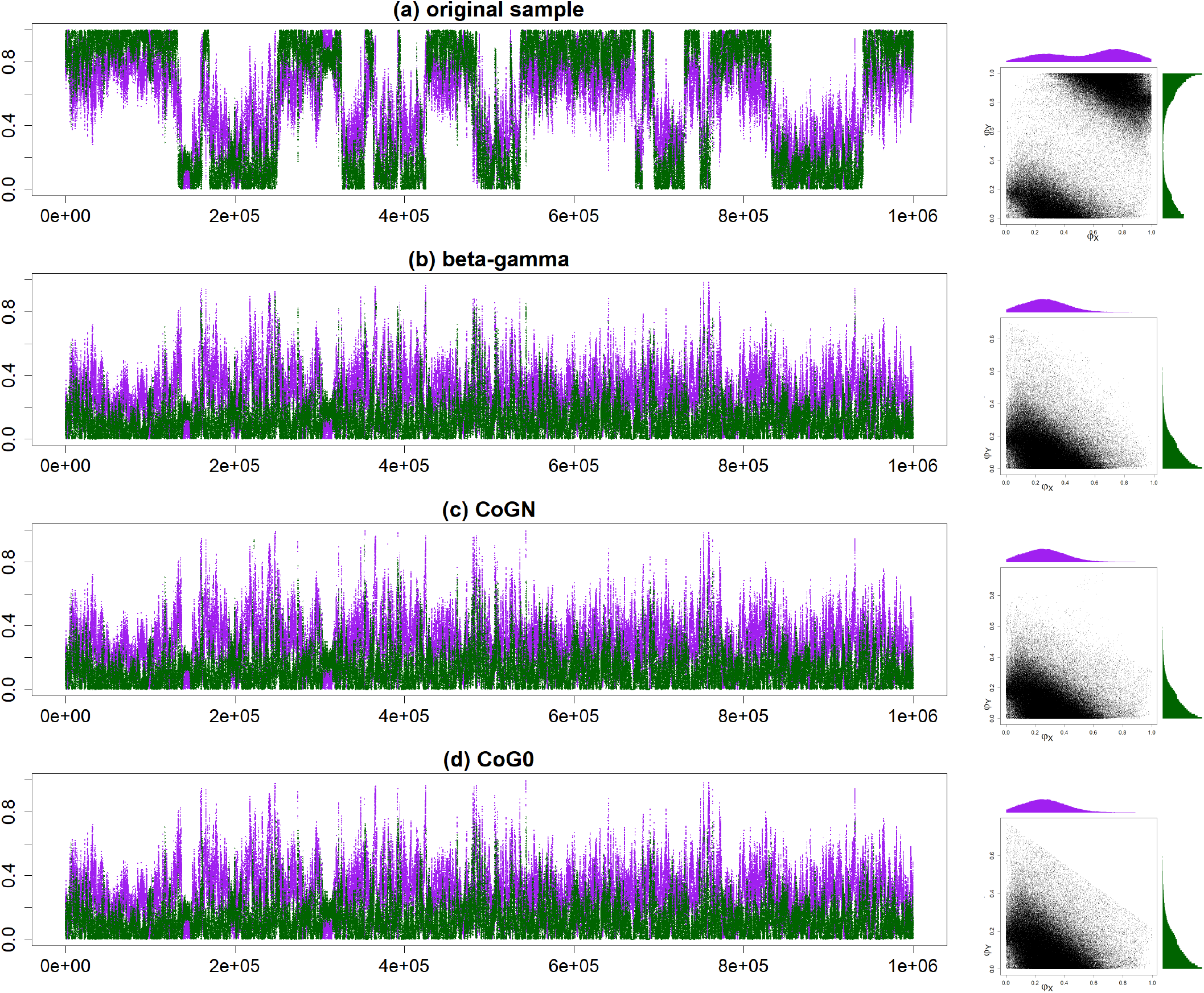
Analysis of the first 500 exonic loci on chromosome 1 from the *Heliconius* data. See legend to figure 3.

**Figure S2:**
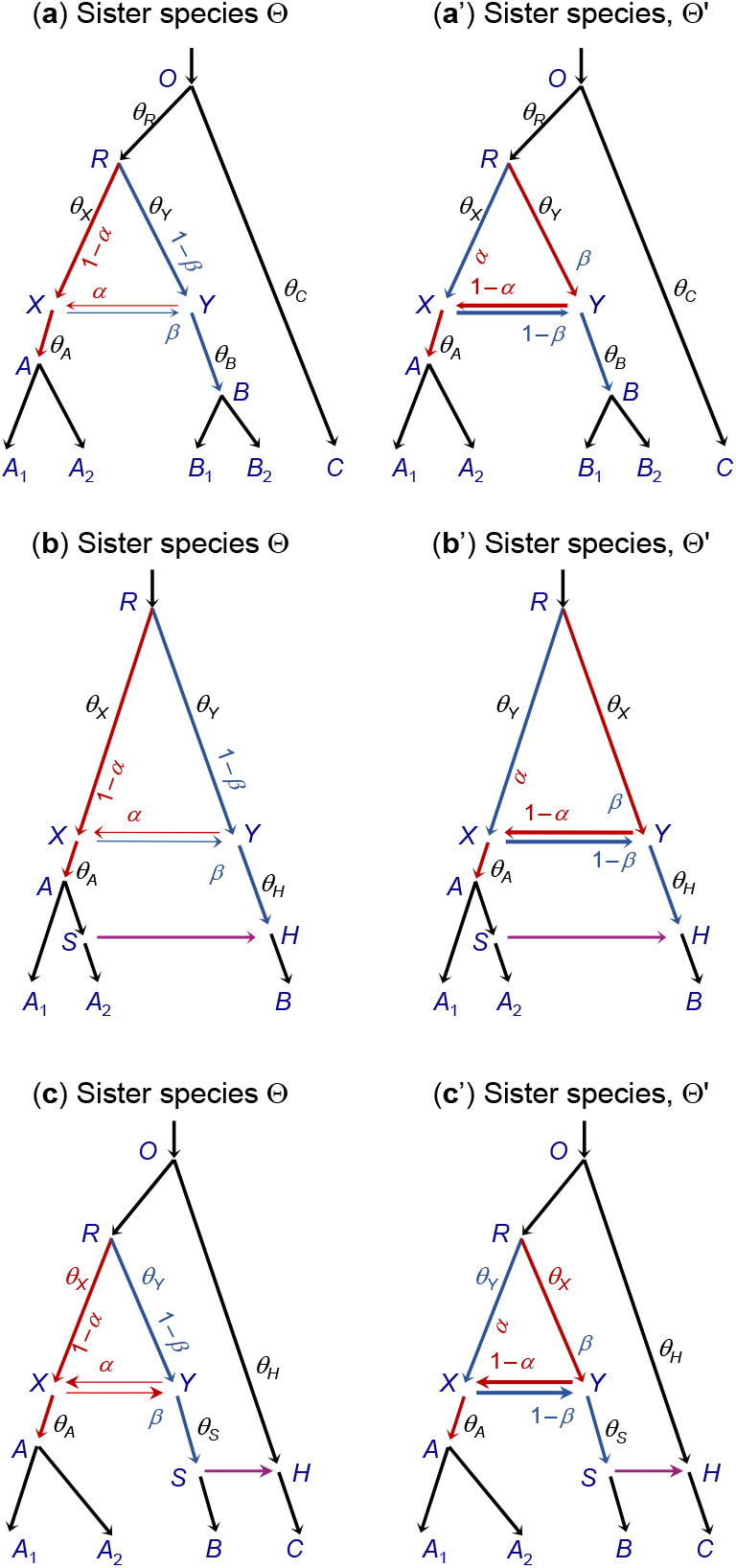
Three species trees (MSci models), each with a BDI event between sister species, exhibiting within-model unidentifiability. (**a & a**′) Subtrees are added to branches *A, B*, and *R* in the basic model of figure 1a. (**b & b**′) A BDI event between sister species *X* and *Y* with a unidirectional introgression involving descendant branches of *X* and *Y*. (**c** and **c**′) A BDI event between sister species *X* and *Y* with a unidirectional introgression involving one descendant branch and another branch that is not a descendant of *X* or *Y*. In all three cases, the parameter mapping is 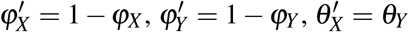, and 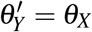.

**Figure S3:**
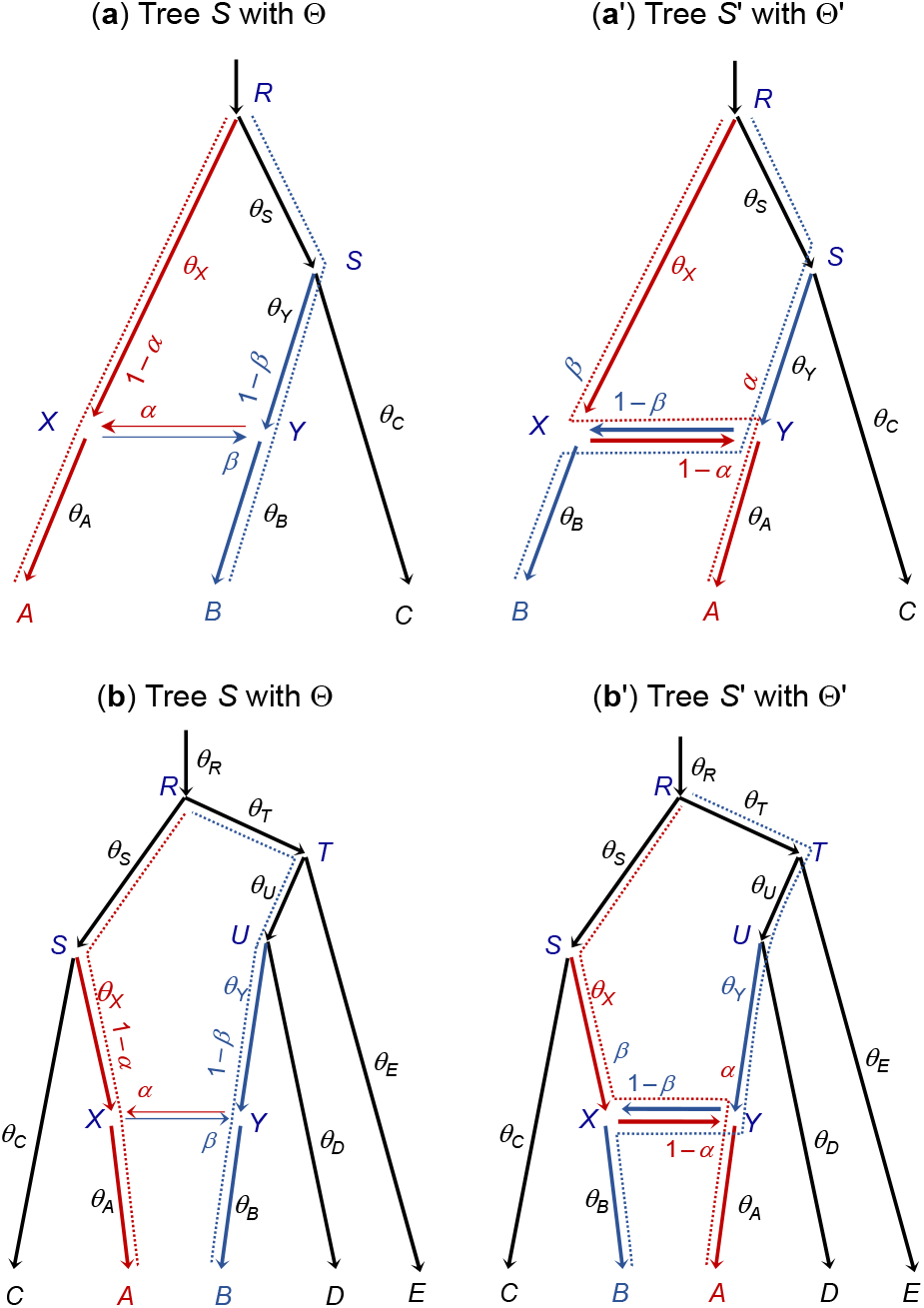
Two pairs of species trees or unidentifiable MSci models with a BDI event between non-sister species creating cross-model unidentiability. (**a & a**′) A pair of unidentifiable models with a BDI event between non-sister species. The dotted lines indicate the main routes taken by sequences sampled from species *A* and *B*, if the introgression probabilities *α* and *β* are 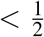. (**b & b**′) Another pair of unidentifiable models with a BDI event between non-sister species. The parameter mapping from Θ to Θ′ in both cases is 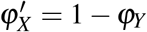 and 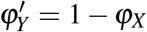, with all other parameters (such as *θ*_*X*_, *θ*_*Y*_, *θ*_*A*_, and *θ*_*B*_) to be identical between Θ and Θ′.

**Figure S4:**
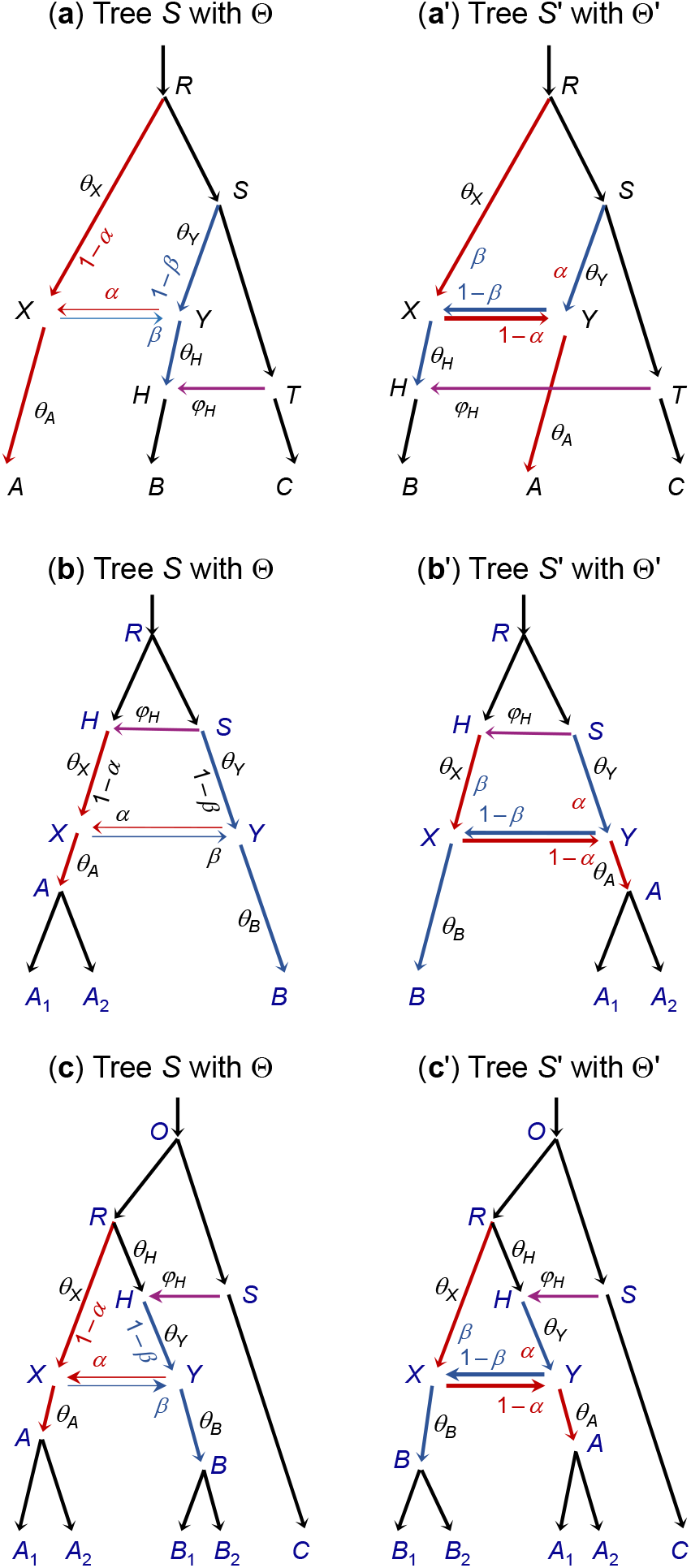
Three pairs of species trees (or unidentifiable MSci models) with one BDI event between non-sister species, illustrating the mapping of parameters (Θ and Θ′). In (**a**), *RXA* and *SYH* are non-sister species. In (**b**) & (**c**), nodes *X* and *Y* are non-sister species because of the unidirectional introgression event involving branches *RX* and/or *RY*. In each of the three cases, the mirror model (*S*′ with Θ′) is generated by pruning off branches *AX* at *X* and *BY* at *Y*, swapping places and reattaching, and applying the mapping 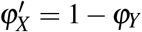 and 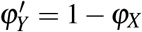.

**Figure S5:**
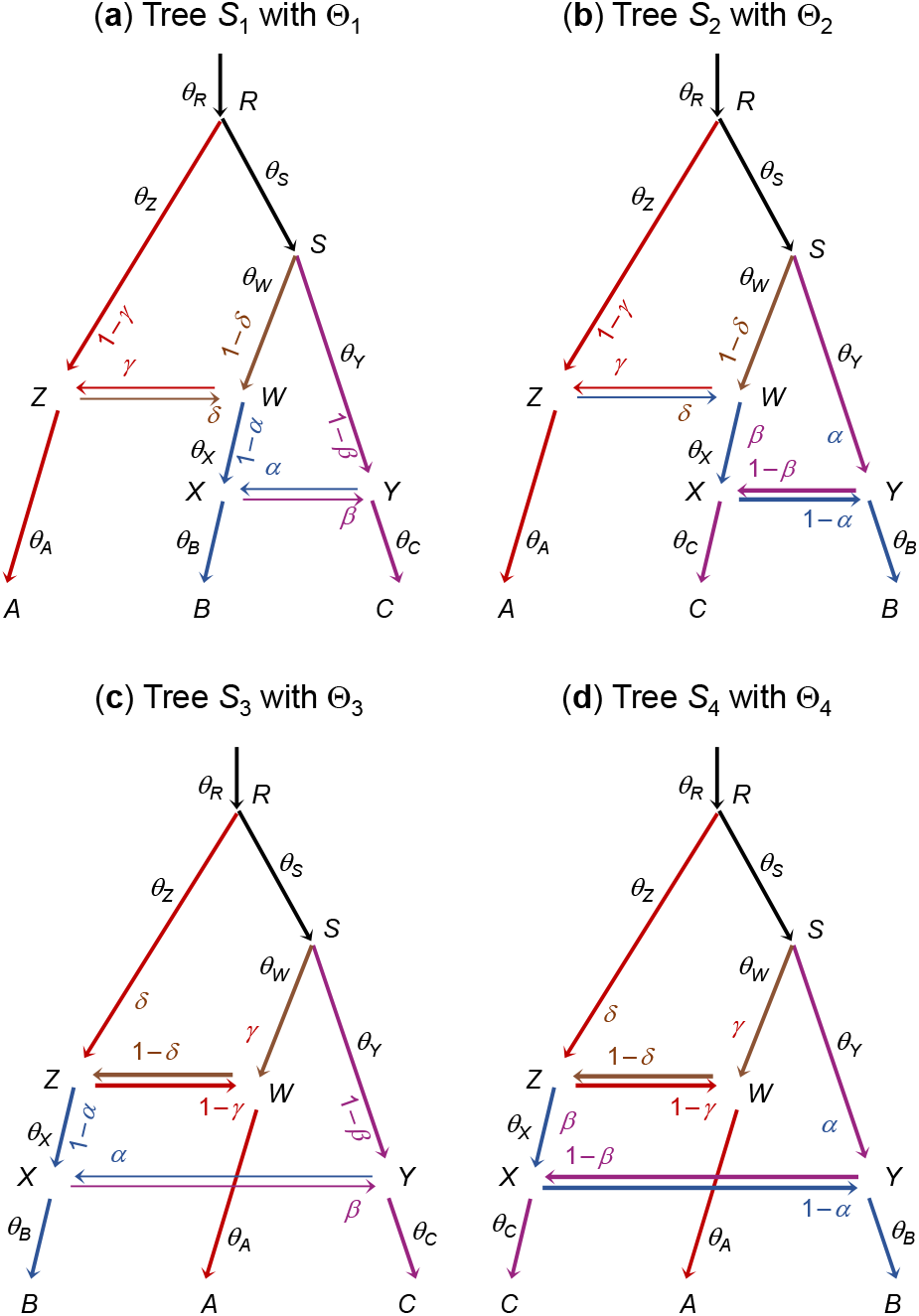
Four species trees for species *A, B*, and *C* representing four unidentifiable models each with two BDI events between non-sister species. The cross-model parameter mappings concern only the introgression probabilities *φ*_*X*_ ≡ *α, ϕ*_*Y*_ ≡ *β, φ*_*Z*_ ≡ *γ*, and *φ*_*W*_ ≡ *δ*, while all other parameters are the same among the models. The colored lines indicate the main routes taken by sequences sampled from *A* (red), *B* (blue), and *C* (purple), if the introgression probabilities *α, β, γ*, and *δ* are all 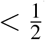, from which the unidentifiability of the four models can be seen easily. Based on figure S9 of Finger *et al*. (2021).

**Figure S6:**
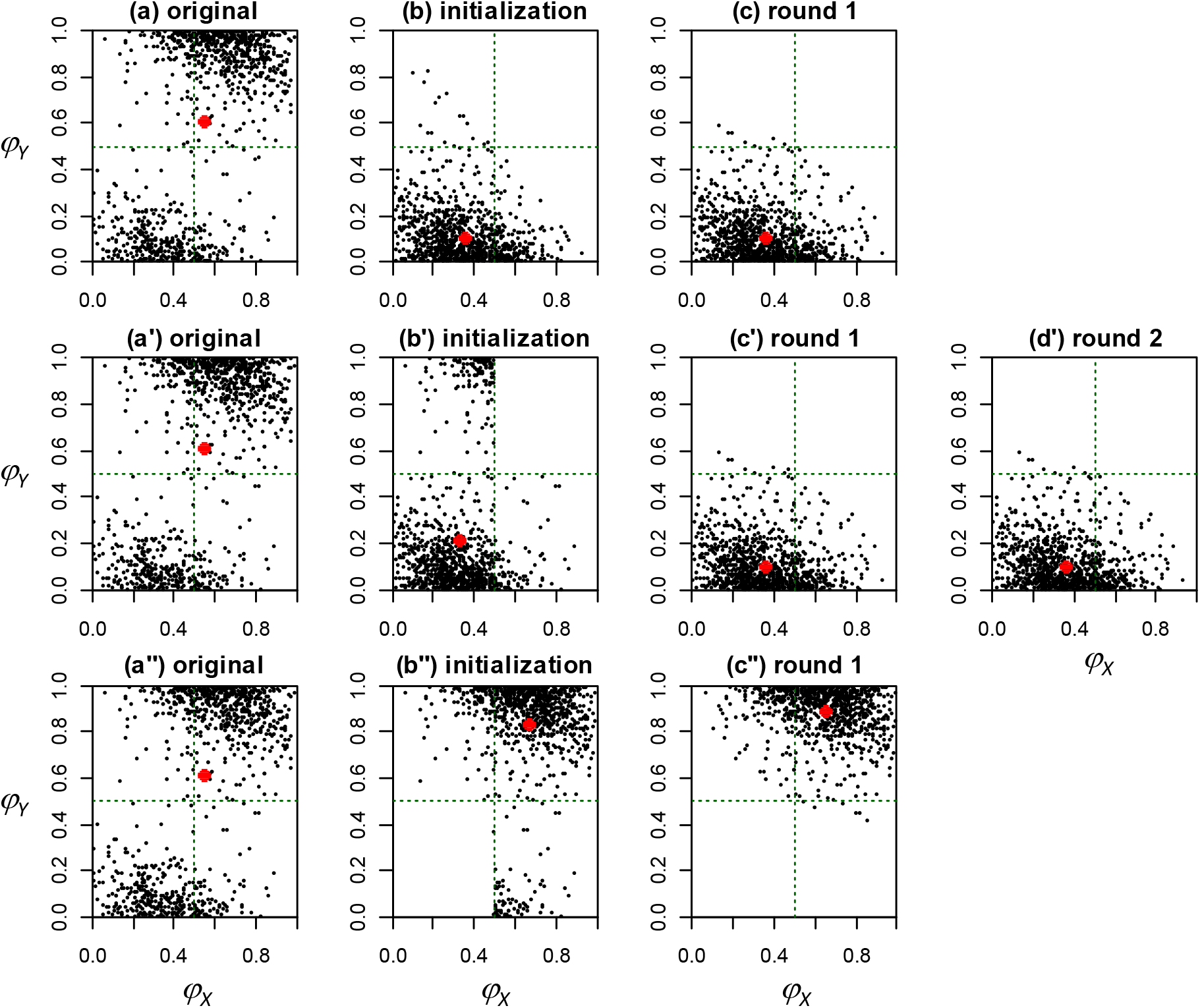
The CoGN_0_ algorithm moves sampled points to their mirror positions to be as close as possible to the center of gravity. Note that (*φ*_*X*_, *φ*_*Y*_) and its mirror position (1 − *φ*_*X*_, 1 − *φ*_*Y*_) are mirror reflections of each other around the point 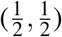. The ‘original’ sample consists of 1000 points, obtained from ‘thinning’ the MCMC sample from the BPP analysis of the 500 noncoding *Heliconius* loci of figure 3a. The mean (*φ*_*X*_, *φ*_*Y*_) = (0.544, 0.614) is indicated by the red dot in **a, a**^′ &^ **a**^′′^. The three rows illustrate three runs of the CoGN_0_ algorithm with different starting positions: (**a**-**c**) *φ*_*X*_ + *φ*_*Y*_ *<* 1, (**a**′-**d**′) 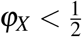 or 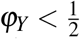, and (**a**″-**c**″) 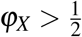 or 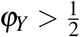. In the first run, the initialization (under the condition *φ*_*X*_ + *φ*_*Y*_ *<* 1) moves 647 points above or right of the line *φ*_*X*_ + *φ*_*Y*_ = 1 to their mirror points below or left of the line, with the new mean (0.348, 0.111), indicated by the red dot (**b**). The algorithm then attempts to move points to their mirror positions to be closer to the red dot. Ten such points are moved, with the new mean (0.353, 0.107) (**c**). In the next iteration, no points move, so the algorithm terminates. In the second run (**a**′-**d**′), the initialization (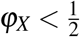 or 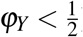) moves 512 points from the upper right corner to their mirror points in the lower bottom, with the new mean (0.327, 0.216) (**b**′). Round 1 moves 136 points, with the new mean (0.353, 0.107) (**c**′). Round 2 moves one point, with the new mean (0.353, 0.107) (**d**′). In the third run, the initialization (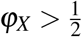 or 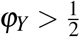) moves 275 points from the lower bottom corner to their mirror points in the upper right corner, with the new mean (0.662, 0.832) (**b**^′′^). Round 1 moves 75 points, with the new mean (0.647, 0.892), and the next round does not move any points, so the algorithm ends. Note that the first two runs converge to the same mean (0.353, 0.107), while the third run converges to its mirror point (0.647, 0.892). If the original positions are taken as the initial positions (i.e., without initialization), the algorithm converges, after one iteration, to (0.647, 0.892), as in the second run. Note that the algorithm operates on four parameters Θ = (*φ*_*X*_, *φ*_*Y*_, *θ*_*X*_, *θ*_*Y*_) but only (*φ*_*X*_, *φ*_*Y*_) is shown here.

**Figure S7:**
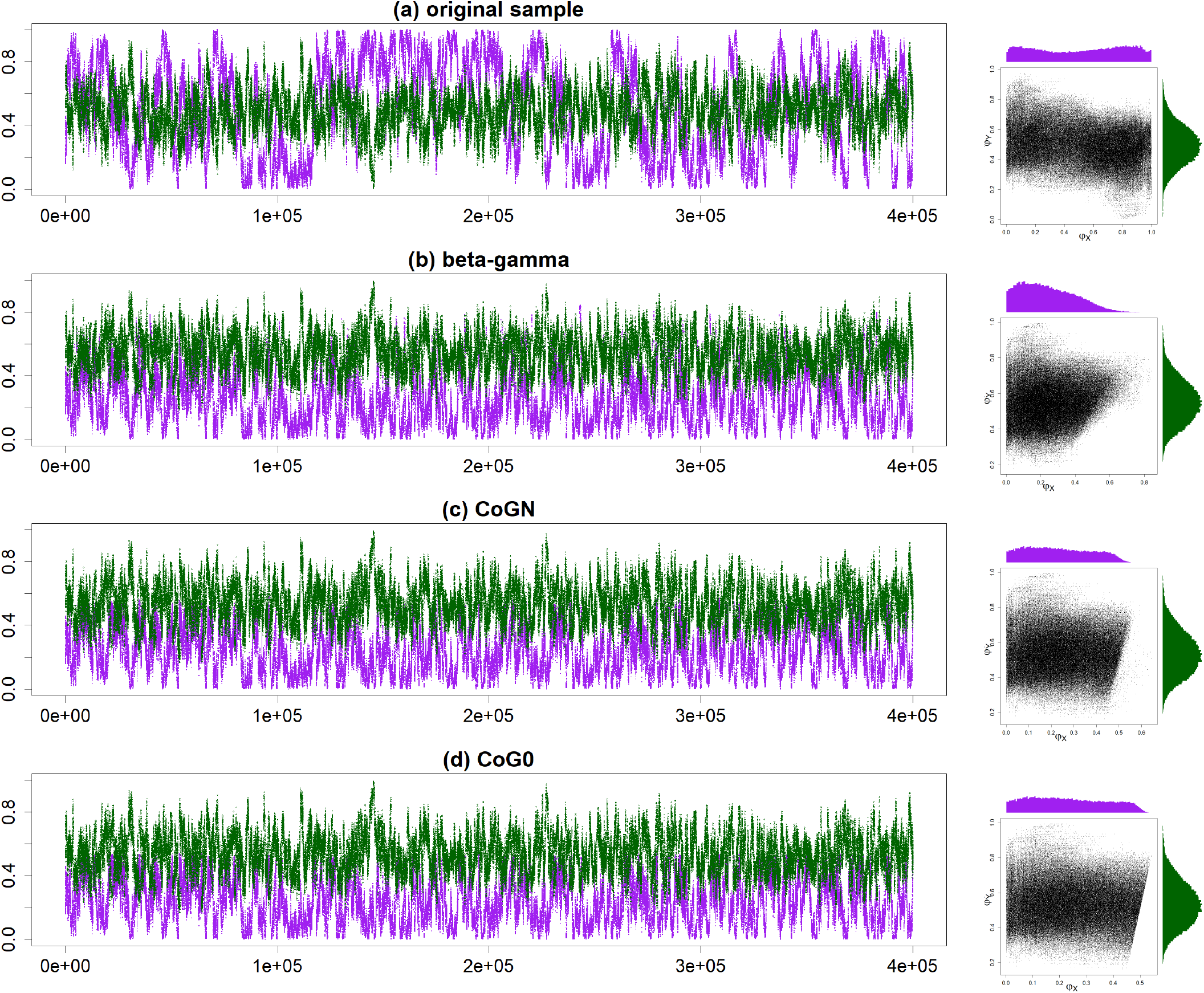
Trace plots of MCMC samples for *φ*_*X*_ (purple) and *φ*_*Y*_ (green) and 2-D scatter plots from BPP analysis of a dataset of *L* = 500 loci simulated under the BDI model of figure 1a. See table S1 for the true parameter values and posterior summaries. The plots are for, from top to bottom, (**a**) unprocessed sample and processed samples using (**b**) the *β*−*γ*, (**c**) the CoG_*N*_, and (**d**) the CoG_0_ algorithms. The true parameter values are Θ = (*φ*_*X*_, *φ*_*Y*_) = (0.7, 0.2), and the post-processing using all three algorithms mapped the samples to the mirror tower around Θ′ = (0.3, 0.8).

**Table S1.** Posterior means and 95% HPD CIs (in parenthees) for parameters in the MSci model of figure 1a from two simulated datasets of *L* = 500 and 1000 loci.

| | truth ($\Theta$) | mirror ($\Theta'$) | beta-gamma | CoG<sub>N</sub> | CoG<sub>0</sub> |
| --- | --- | --- | --- | --- | --- |
| $L = 500$ loci | | | | | |
| $\tau_R$ | 0.01 | | 0.0098 (0.0088, 0.0108) | | |
| $\tau_X = \tau_Y$ | 0.005 | | 0.0050 (0.0045, 0.0055) | | |
| $\theta_A$ | 0.002 | | 0.0020 (0.0018, 0.0021) | | |
| $\theta_B$ | 0.01 | | 0.0101 (0.0093, 0.0108) | | |
| $\theta_R$ | 0.002 | | 0.0020 (0.0006, 0.0034) | | |
| $\theta_X$ | 0.002 | 0.01 | 0.0063 (0.0005, 0.0130) | 0.0066 (0.0005, 0.0133) | 0.0066 (0.0005, 0.0133) |
| $\theta_Y$ | 0.01 | 0.002 | 0.0071 (0.0022, 0.0124) | 0.0067 (0.0017, 0.0120) | 0.0068 (0.0017, 0.0121) |
| $\phi_X$ | 0.7 | 0.3 | 0.755 (0.472, 0.999) | 0.764 (0.528, 0.999) | 0.765 (0.530, 0.999) |
| $\phi_Y$ | 0.2 | 0.8 | 0.447 (0.209, 0.670) | 0.461 (0.214, 0.695) | 0.462 (0.212, 0.695) |
| $L = 2000$ loci | | | | | |
| $\tau_R$ | 0.01 | | 0.0101 (0.0094, 0.0108) | | |
| $\tau_X = \tau_Y$ | 0.005 | | 0.0051 (0.0048, 0.0054) | | |
| $\theta_A$ | 0.002 | | 0.0020 (0.0019, 0.0021) | | |
| $\theta_B$ | 0.01 | | 0.0100 (0.0097, 0.0104) | | |
| $\theta_R$ | 0.002 | | 0.0018 (0.0009, 0.0027) | | |
| $\theta_X$ | 0.002 | 0.01 | 0.0037 (0.0008, 0.0062) | 0.0037 (0.0009, 0.0062) | 0.0050 (0.0006, 0.0097) |
| $\theta_Y$ | 0.01 | 0.002 | 0.0076 (0.0049, 0.0108) | 0.0076 (0.0048, 0.0108) | 0.0064 (0.0019, 0.0104) |
| $\phi_X$ | 0.7 | 0.3 | 0.545 (0.178, 0.903) | 0.545 (0.178, 0.903) | 0.656 (0.449, 0.887) |
| $\phi_Y$ | 0.2 | 0.8 | 0.398 (0.198, 0.598) | 0.398 (0.198, 0.598) | 0.450 (0.205, 0.684) |
| $L = 8000$ loci | | | | | |
| $\tau_R$ | 0.01 | | 0.0098 (0.0094, 0.0102) | | |
| $\tau_X = \tau_Y$ | 0.005 | | 0.0049 (0.0048, 0.0051) | | |
| $\theta_A$ | 0.002 | | 0.0020 (0.0019, 0.0020) | | |
| $\theta_B$ | 0.01 | | 0.0100 (0.0098, 0.0102) | | |
| $\theta_R$ | 0.002 | | 0.0021 (0.0017, 0.0025) | | |
| $\theta_X$ | 0.002 | 0.01 | 0.0045 (0.0003, 0.0083) | 0.0044 (0.0003, 0.0081) | 0.0051 (0.0003, 0.0106) |
| $\theta_Y$ | 0.01 | 0.002 | 0.0095 (0.0059, 0.0129) | 0.0096 (0.0060, 0.0130) | 0.0089 (0.0041, 0.0128) |
| $\phi_X$ | 0.7 | 0.3 | 0.645 (0.442, 0.848) | 0.645 (0.436, 0.848) | 0.645 (0.436, 0.846) |
| $\phi_Y$ | 0.2 | 0.8 | 0.334 (0.128, 0.632) | 0.336 (0.129, 0.639) | 0.310 (0.123, 0.539) |

Note.— Empty values for $\Theta'$ mean the same values as for $\Theta$ . MCMC samples are processed using the three algorithms and then summarized. See figure S7 for the tracecatter plots for the dataset of $L = 500$ . The datasets, with each locus consisting of four sequences per species (or eight sequences per locus) and 500 sites per sequence, are simulated using the true parameter values ( $\Theta$ ).

